# Phenotypic and genomic differentiation of *Arabidopsis thaliana* along altitudinal gradients in the North Italian alps

**DOI:** 10.1101/023051

**Authors:** Torsten Günther, Christian Lampei, Ivan Barilar, Karl J. Schmid

## Abstract

Altitudinal gradients represent short-range clines of environmental parameters like temperature, radiation, seasonality and pathogen abundance, which allows to study the footprints of natural selection in geographically close populations. We investigated phenotypic variation for frost resistance and light response in five *Arabidopsis thaliana* populations ranging from 580 to 2,350 meters altitude at two different valleys in the North Italian Alps. All populations were resequenced as pools and we used a Bayesian method to detect correlations between allele frequencies and altitude while accounting for sampling, pooled sequencing and the expected amount of shared drift among populations. The among population variation to frost resistance was not correlated with altitude. An anthocyanin deficiency causing a high leaf mortality was present in the highest population, which may be non-adaptive and potentially deleterious phenotypic variation. The genomic analysis revealed that the two high-altitude populations are more closely related than the geographically close low-altitude populations. A correlation of genetic variation with altitude revealed an enrichment of highly differentiated SNPs located in genes that are associated with biological processes like response to stress and light. We further identified regions with long blocks of presence absence variation suggesting a sweep-like pattern across populations. Our analysis indicate a complex interplay of local adaptation and a demographic history that was influenced by glaciation cycles and/or rapid seed dispersal by animals or other forces.

## Introduction

Local adaptation results from interactions of populations with biotic and abiotic environmental conditions. Numerous phenotypic traits that contribute to adaptation show therefore clinal variation along environmental gradients (Huxley, 1938). Phenotypic clines have been investigated for many years (Mayr, 1942) because predictions regarding patterns and processes of natural selection can be tested in a rigorous fashion (Etterson & Shaw, 2001). In annual plants, an example of a well studied cline is the change of flowering time with latitude gradients (Stinchcombe *et al*, 2004). The timing of the transition from the vegetative to the reproductive stage is strongly related to fitness in annual plants and therefore depends on day length and temperature.

In mountaineous environments, altitude is one of the strongest environmental gradients because abiotic parameters like temperature, radiation intensity, atmospheric pressure, and partial pressure of gases simultaneously change over short distances (Körner, 2007). The biotic environment includes competitors, herbivores, pathogens or symbionts whose occurrence differs with altitude as well. The combination of these factors results in different ecological niches at different altitudes, which likely exert a strong selection pressure for local adaptation. In addition to external factors, the intrinsic genetic potential for adaptation is affected by altitude since the absolute area available for plant growth tends to rapidly decrease with altitude (Körner, 2007) causing small population sizes and increasing levels of genetic drift and inbreeding at high altitudes (Thiel-Egenter *et al*, 2011; Manel *et al*, 2012).

Another critical factor in adaptation to different altitudes is the age of local populations. Long-term changes in temperature and other environmental parameters caused by glacial cycles contributed to large fluctuations in the altitudinal distribution of species, which retreated from high altitudes during glacial maxima and expanded during interglacial periods. However, numerous stable glacial refugial populations on mountain tops (nunataks) existed in the European Alps (Schönswetter *et al*, 2005) and other mountain ranges. In such populations, selection and genetic drift likely contributed to the rapid genetic and phenotypic divergence from other populations of the same species, and frequently led to the formation of endemic sibling species restricted to high altitudes.

*Arabidopsis thaliana* shows a wide distribution and local adaptation within the Northern hemisphere for many morphological, life history and other fitness-related traits (reviewed by Koornneef *et al*, 2004; Bergelson & Roux, 2010). Genomic surveys suggested adaptation to large-scale environmental gradients in central Europe (Clark *et al*, 2007; Hancock *et al*, 2011; Horton *et al*, 2012), the Iberian Peninsula (Méndez-Vigo *et al*, 2011) and Scandinavia (Long *et al*, 2013; Huber *et al*, 2014). Adaptation to different altitudes can also be studied in *A. thaliana* since it occurs from the sea level up to 4,250 m altitude (Al-Shehbaz & O’Kane, 2002). High altitude adaptation was observed in populations in the eastern Pyrenees and included traits like fecundity, phenology and biomass allocation, suggesting selection for higher vigour in high altitude plants (Montesinos-Navarro *et al*, 2011), selection for alpine dwarfism in the Swiss Alps (Luo *et al*, 2015), or a constitutive protection against UV-B radiation protection in Shakdara, Central Asia at 3,063 m a.s.l (Biswas & Jansen, 2012).

We investigated the roles of adaptive evolution and demographic history in the distribution of genetic variation and putative adaptive phenotypic traits along an altitudinal gradient in the European Alps. We sampled five *A. thaliana* populations at different altitudes up to 2,300 m in the South Tyrolia and Trento provinces of Italy. Four populations represent two pairs of high and low altitudes in two different valleys, and and a fifth population was located at equal distance to both gradients. In these populations, we characterized phenotypic differentiation in frost resistance and response to light and UV-B stress because these environmental parameters are strongly associated with altitude (Blumthaler *et al*, 1997; Körner, 2007). Accessions from the Northern Italian Alps were previously shown to be genetically distinct from other regions and to harbour reduced genetic diversity (Cao *et al*, 2011). We reasoned that high altitude adaptation resulted in a strong signal of genetic differentiation relative to the genome-wide background, which should allow to identify selected genomic regions. We sequenced pools of individuals from each of the five populations, because pool sequencing is a cost-efficient means to analyse diversity in populations to infer demography and selection, and was previously applied in several plant and animal species (e.g. Turner *et al*, 2010; He *et al*, 2011; Kolaczkowski *et al*, 2011; Fabian *et al*, 2012; Lamichhaney *et al*, 2012; Orozco-terWengel *et al*, 2012; Fischer *et al*, 2013). Although sequencing of pools comes at the cost of noisy estimates and a loss of linkage information (Futschik & Schlötterer, 2010; Zhu *et al*, 2012), suitable frameworks that account for the special properties of pooled sequence data are available (Futschik & Schlötterer, 2010; Kofler *et al*, 2011b,a; Boitard *et al*, 2012; Günther & Coop, 2013). We investigated the demographic history by Approximate Bayesian Computation (ABC, Beaumont *et al*, 2002), and correlated allele frequencies with altitude to identify genomic regions that are significantly differentiated between high and low altitude populations because of local adaptation. We observed a complex pattern of genomic and phenotypic diversity suggesting that adaptation to high altitudes may affect genes involved in light responses and soil conditions as well as other stress factors, but also indicate that there is no simple pattern of genomic and phenotypic differentiation with altitude.

## Materials and Methods

### Plant material

Seeds were sampled at five locations in the alps of South Tyrolia and Trento provinces to represent two pairs of high and low altitudes, respectively, in different valleys and mountain ranges but in close geographic proximity (Table 1 and Figure 1). All populations were located on steep slopes and exposed to the South (Coordinates: Vioz/Coro 46°22’57”N, 10°39’35”E; Terz 46°21’50”N, 10°55’13”E; Finail 46°44’31”N, 10°49’5”E; Juval 46°38’55”N, 10°58’27.85”E; Laatsch 46°40’18.00”N, 10°30’54.00”E). Two low altitude populations (Laatsch and Juval) are located on scree which was partly overgrown in Juval, but still moving in Laatsch. These two sites were the least disturbed of the five sites. The third valley population (Terz) is located on marginal soils in dry stone walls in the vicinity of a wine yard and was highly disturbed. The two high-altitude populations occur at South exposed rocky outcrops that serve as resting places of mountain goats, and therefore are strongly disturbed habitats with a high supply of nitrogen and phosphorous. Due to different seasonality, we visited the low altitude populations in early May and the high altitude populations in mid-July. All locations were sampled in a spatially balanced design to include the maximal diversity present at the site. Plants were cultivated for DNA extraction on standard gardening soil (Einheitserde ED 63T). Forty day old whole plants were freeze dried in a Christ Alpha 14 (Martin Christ Gefriertrocknungsanlagen GmbH, Osterode am Harz, Germany) freeze dryer before DNA extraction.

**Table 1:** Summary of pooled sequencing libraries.

| Population | Individuals<br>sequenced | Altitude [m] | PoolSeq summary statistics |  |  |  |
| --- | --- | --- | --- | --- | --- | --- |
| | | | Median coverage<br>of nDNA | Percent<br>genome covered | Nucleotide<br>diversity, $\pi$ (SE) | Tajima's $D$ (SE) |
| Juval | 29 | 577 - 657 | 32x | 94.3 | 0.00305 (0.00003) | 0.0352 (0.0063) |
| Terz | 25 | 810 - 945 | 31x | 95.8 | 0.00439 (0.00003) | -0.0107 (0.004) |
| Laatsch | 29 | 970 - 1010 | 19x | 94.5 | 0.00190 (0.00003) | -0.1912 (0.0084) |
| Finail | 29 | 2250 - 2270 | 32x | 94.2 | 0.00190 (0.00002) | -0.7567 (0.0046) |
| Vioz/Coro | 19 | 2210 - 2355 | 50x | 93.3 | 0.00166 (0.00002) | -0.7122 (0.0044) |

**Figure 1:**
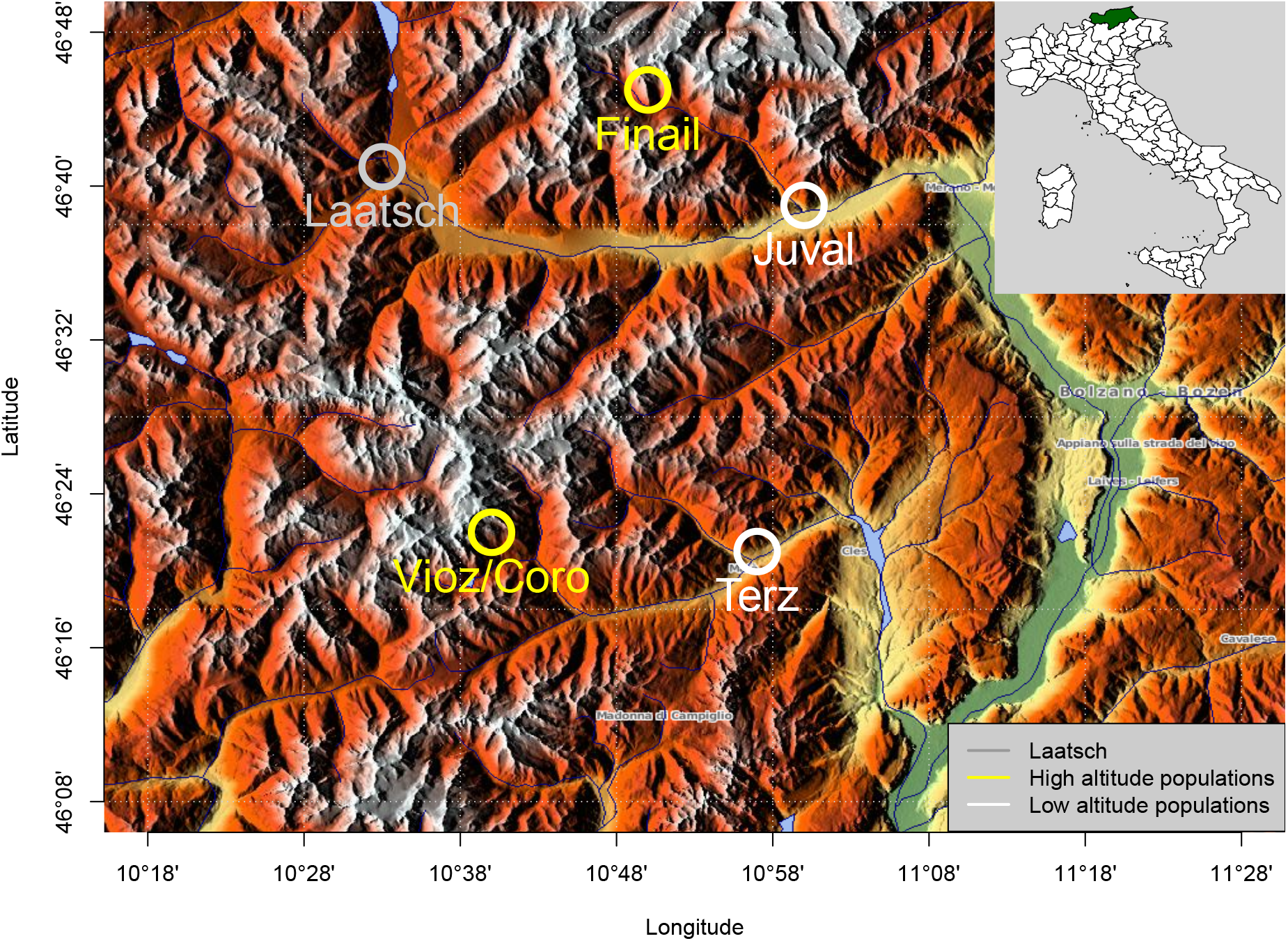
Geographic map showing the location of the five populations included in this study. Map source: http://www.maps-for-free.com/

### Phenotypic analysis of spring freezing tolerance and response to light stress and UV-B radiation

#### Freezing tolerance

We investigated the plants for differences in freezing damage on the population level under two temperature regimes with plants from all populations grown in the greenhouse by adapting the method of Martin *et al* (2010). Eight random accessions per population were tested at a minimum temperature of -15°C, and five random accessions per population at -7°C minimum temperature. We raised the plants under long day condition (16 h light/8 h dark) at 23° C and adapted them to the cold for one day at 4° C at the eight leaf stage. Whole plants were sampled, washed with deionized water and wrapped in aluminum foil after removal of residual water.

We then put the samples into a steel vacuum Dewar bottle which was placed in a styrofoam box (14 l volume and 5 cm wall thickness). Two temperature loggers (3M™Temperature Logger TL20, 3M Deutschland GmbH, Neuss) were added to the Dewar bottle above and below the samples to monitor the realized treatment temperature. To ensure a slow temperature decrease, we added a PET-bottle with 1 l of saturated salt water (20% NaCl solution) to the styrofoam box. The box was kept at -20° C for 17 and 7 hours, respectively. Slow defrosting was ensured at 4°C for approximately 20 h. Samples were put into 10 ml of deionized water in sterile 15 ml centrifuge tubes and left at room temperature overnight to allow cytosol leakage of damaged cells. We then measured electric conductivity with a handheld conductivity meter (WTW LF 90, KLE1 sensor; measure 1). To obtain conductivity at 100% cell damage, we autoclaved all samples (Systec DB-23 benchtop autoclave) and measured conductivity again (measure 2). Freezing damage of each sample was calculated as the proportion of measure 1 over measure 2.

#### Light and UV-B stress

We further characterized the response to different light conditions. Random individuals of each population were cultivated on standard gardening soil (Einheitserde ED 63T) at 23c in a climate cabinet (GroBank BB-XXL^3+^, CLF Plant Climatics, Emersacker, Germany) under three light treatments under long day conditions (16/8 light/dark): (1) A control low light treatment, (2) a low light with UV-B treatment and (3) a light-stress treatment without UV-B. The low light treatment consisted of 168 *μ*mol m^-2^ s^-1^ fluorescent light (Philips Master TL-D 58W/840 Reflex), and the UV-B treatment used the same fluorescent setting with additional UV-B radiation exposure three times per day for one hour at variable intensities (6:00*-*7:00 am 0.01 mWcm^*-*2^, 11:00*-*12:00 am 0.027 mWcm^*-*2^ and 4:00-5:00 pm 0.027 mWcm^*-*2^). In the light-stress treatment, we exposed plants to fluorescent light with an intensity of 470 *μ*mol m^*-*2^ s^*-*1^, which is more than twice the optimal light intensity for the Col-0 ecotype (200 *μ*mol m^*-*2^ s^*-*1^). In the UV-B treatment, fluctuating UV-B levels were judged to be more realistic than stable UV-B exposure because frequent cloud cover reduces the total annual amount of light and UV-B radiation but fluctuations in light intensity increase strongly with higher altitude (Körner, 2007). The leaf color of each plant was used as stress indicator because light or UV-B stressed plants show increased anthocyanin production which causes a red leaf colour (Chalker-Scott, 1999). All plants were photographed with a digital camera (Samsung NX 10) at high resolution from the top and customized macros of the ImageJ distribution Fiji (Schindelin *et al*, 2012) were used to measure the red leaf area, the total leaf area and the area of dead plant material.

#### Statistical analysis of phenotypic data

To identify differences in freezing damage between populations, we fitted a generalized linear model (GLM) with freezing damage odds as dependent variable and a population factor and a test-condition factor and their interaction as independent variables based on a quasibinomial error distribution (logit link) as implemented in the R package 

~~~
stats
~~~

 (DevelopmentCoreTeam, 2014). Although statistical theory suggests to use a binomial error distribution to analyse the proportion of read and dead tissue after the light treatment, we used a general least squares model (GLS) assuming a normal distributed error because the residual distribution did not deviate from a normal distribution (Shapiro-Wilk test, W = 0.96, p = 0.19) and only the GLS model allows to correct for the strong heteroscedasticity in the data. Using the 

~~~
nlme
~~~

 package in R (Pinheiro *et al*, 2014) we fitted GLS models with the red proportion or the dead proportion as dependent variable and population, treatment and their interaction as independent factors. We allowed for different variances for each level of population and treatment using the *varComb* function with two *varIdent* functions. To identify differences between treatments in both experiments, we performed *post-hoc* comparisons with the *glht* function from the R package 

~~~
multcomp
~~~

 and corrected for the false discovery rate (FDR; Benjamini & Hochberg, 1995).

### Sequencing and population genetic analysis

#### DNA sequencing

We ground dry whole plants with a Retsch MM400 centrifugal mill (Retsch GmbH, Haan, Germany) and extracted the DNA with a standard CTAB maxi prep protocol. DNA quality and concentration were determined on a 0.8% agarose gel and additionally with the Qubit 2.0 Fluorometer. For each of the five populations, we prepared a pooled DNA sample (5 *μ*g) by mixing of equimolar amounts of DNA from all individuals. Each DNA pool was then converted into a barcode-tagged genomic sequencing library using the standard Illumina protocol. The five tagged libraries were pooled and sequenced together on a single lane of an Illumina HiSeq 2000 sequencer by GATC Biotech (Konstanz, Germany). We then trimmed the 101 bp long paired-end reads to consecutive runs of base calls with a quality *≥* 25 and filtered for a minimum read length of 35 bp using SolexaQA (Cox *et al*, 2010). The trimmed high-quality reads were mapped against the *A. thaliana* TAIR10 reference genome (Lamesch *et al*, 2012) using BWA version 0.6.2 (Li & Durbin, 2009) using the non-default parameters -n 0.05 (mismatch rate) and disabled seeding of reads. The BWA module 

~~~
sampe
~~~

 mapped paired-end reads to a single SAM file and samtools version 0.1.18 (Li *et al*, 2009) filtered the resulting alignments to include only properly paired reads and a minimum mapping quality of 20. For all downstream analyses, we converted the data to pileup files with samtools 

~~~
mpileup
~~~

. We then identified SNPs as polymorphic sites of at least 12x coverage in at least one pool with a minimum of two reads supporting a non-reference allele and polarized them with the close relative *Arabidopsis lyrata* (Hu *et al*, 2011).

#### Population genomic analysis

We further analysed the Pileup files with Popoolation version 1.2.2 (Kofler *et al*, 2011a) to calculate nucleotide diversity, *π*, Wattersons *θ* and Tajima’s *D* separately for each population. First, each position was independently subsampled to create a uniform 18x coverage over all populations by random drawing base calls from the pileup file without replacement. Then, *π*, *θ* and Tajima’s *D* were calculated separately for each population using sliding windows with window and step sizes of 2,000 bp each. Only sites with a coverage of at least 10 and maximally 100 were included in the next analysis steps.

We annotated SNPs with SnpEff version 3.1 (Cingolani *et al*, 2012) based on the TAIR10 annotation (Lamesch *et al*, 2012) and assigned them to one of the following classes: intergenic, 5’ UTR, 3’ UTR, intron, synonymous coding, non-synonymous coding, synonymous stop, stop lost, stop gained and start lost. SNPs belonging to different groups (e.g. non-synonymous coding and stop gained) were assigned to the stronger effect category (e.g., stop gained). We further annotated regulatory SNPs based on known and predicted transcription factor binding sites from AGRIS (Yilmaz *et al*, 2011). The effect of non-synonymous SNPs on protein function (Günther & Schmid, 2010) was determined with the MAPP program (Stone & Sidow, 2005), which classifies nonsynonymous SNPs into deleterious or tolerated amino acid substitutions based on a collection of homologous proteins obtained with PSI-BLAST searches (Altschul *et al*, 1997) of all *A. thaliana* proteins against Uniprot (Wu *et al*, 2006). We calculated a phylogenetic tree for these proteins with semphy (Friedman *et al*, 2002) and used the multiple protein sequence alignment and the tree as input for MAPP (Stone & Sidow, 2005).

#### Identification of strongly differentiated genes

To identify strongly differentiated genes, we correlated altitude with allele frequencies within populations with Bayenv2.0 because it models the sampling noise inherent to pooled sequencing and accounts for a shared history of populations (Günther & Coop, 2013). To characterize this relationship, we estimated a covariance matrix with Bayenv2.0 from a subsample of 10,000 high confidence SNPs (i.e. SNPs with a coverage between 30 and 100 in each of the five populations) based on 1,000,000 MCMC iterations. To ensure convergence we estimated a second matrix from another subsample of 10,000 SNPs. For the Bayenv2.0 analysis, we included only SNPs for which both alleles segregated in at least two populations and the per population sequence coverage was between 15 and 100, which resulted in a total of 400,231 SNPs. Using these data we estimated the genotype-environment correlation statistics *Z* and *ρ*(*X, Y’*) (Günther & Coop, 2013) between allele frequencies and altitude per population (normalized to mean 0 and standard deviation 1) estimated during 200,000 MCMC iterations.

To further analyse the functional annotation of highly differentiated genes, we searched for an enrichment of different annotation categories obtained from AraPath (Lai *et al*, 2012), which comprises gene ontology (GO, Ashburner *et al*, 2000), KEGG (Kanehisa *et al*, 2012), AraCyc (Mueller *et al*, 2003) and plant ontology (PO, Cooper *et al*, 2013) annotations for all *A. thaliana* genes, and additionally includes gene sets obtained from extensive literature research (Lai *et al*, 2012). To avoid a biased enrichment analysis, we tested enrichment of annotations using Gowinda (Kofler *et al*, 2012), which accounts for gene length using a permutation approach. We conducted 100,000 permutations and included all annotation categories of at least two genes while all SNPs in a gene and within 2,000 bp distance of a transcript were assigned to the respective gene.

#### Demographic history

We investigated the genetic relationship of populations with Neighbour-joining trees (Saitou & Nei, 1987) based on the matrix of average pairwise *F*_*ST*_ values calculated by PoPoolation2, and from a correlation matrix of allele frequencies derived from the covariance matrix estimated by Bayenv2.0. The trees were plotted with the R package 

~~~
ape
~~~

 (Paradis *et al*, 2004). Additionally, we reconstruced the historical relationship among the five populations using their current genome-wide allele frequencies with TreeMix (Pickrell & Pritchard, 2012). We carried out a run which included all non-singleton SNPs segregating in all five populations with a minimum coverage of 20 per population (1,071,527 SNPs). We bootstrapped each run 1,000 times and carried out a four-population test which tests whether the relationship of the two pairs of high and low populations can be explained by any of the possible tree topologies (Reich *et al*, 2009; Patterson *et al*, 2012) to verify the TreeMix results.

We estimated divergence times between pairs of high and low altitude populations by simulating models of demographic history with msABC (Pavlidis *et al*, 2010) followed by an Approximate Bayesian Computation (ABC) analysis with the 

~~~
abc
~~~

 R package (Csilléry *et al*, 2012). We investigated four different models under a population-split scenario to estimate divergence times between low and high altitude populations (Figure S3). All models assumed a common ancestral population with a founding bottleneck leading to the high altitude population, but models differed in their post-split growth parameters of the high altitude populations: (i) the no-growth model assumed a constant population size after the split, (ii) the step-growth model assumed an instantaneous population size change some time after the split, (iii) the growth model assumed an exponential growth of the population after the split where the growth rate was calculated based on a previously drawn current population size (N_0_; Table S3) and the time of the split, and (iv) the exponential model assumed exponential growth of the population size after the split. We simulated each model 200,000 times with and without bidirectional migration and included them in a model selection step with cross-validation based on selected summary statistics (see below) to select the best-fitting model. Posterior model probabilities were calculated with the *postpr* function of the R 

~~~
abc
~~~

 package using the *mnlogistic* method and a tolerance rate of 0.005. Using the best fitting model, we inferred the posterior distribution of the population split time.

In the simulations we assumed uniform priors for parameters (Table S3), and obtained mutation and recombination rates used in simulations from empirical studies (Kim *et al*, 2007; Ossowski *et al*, 2010; Cao *et al*, 2011) or based them on plausibility arguments (size of the high altitude populations, population growth parameter, time of the split). The ABC analysis included the following summary statistics: mean and variance of Tajima’s *D* and *π*, mean of Watterson’s *θ* and average number of segregating sites for each of the populations; *F*_*ST*_ values and numbers of shared, fixed and private SNPs, which were calculated for each population pair. We calculated summary statistics from 2 kb windows and randomly sampled 1,000 windows from the sequence data. Posterior parameter distributions were estimated by comparing observed and simulated summary statistics using the *abc* function of the R 

~~~
abc
~~~

 package with the *rejection* method and a tolerance rate of 0.01. From the posterior distributions of the best models, we calculated the Highest Posterior Density Interval (HPDI 90) of the estimated mean of the population sizes and the time of their split using the 

~~~
LaplacesDemon
~~~

 R package (Hall, 2013). The HPDI 90 measurement gives the most accurate 90% credible interval (Bayesian equivalent of confidence intervals) of the posterior distributions.

## Results

### Phenotypic differences between high and low altitude populations

In the test for freezing resistance we first decreased the temperature to -15°C with a shoulder at around -7°C, which marks the begin of major tissue freezing (Figure 2B). In the second experiment, temperatures were lowered to only -7° C and no shoulder was observed in the temperature curves (Figure 2D). The two freezing treatments caused strong differences in the degree of tissue damage (Figure 2 A+C, F1*/*59=128, p*<*0.001) with a population median from 0.8 to 0.9 in the first test and from 0.1 to 0.6 in the second test. Based on the GLM there was no significant temperature × population interaction (F_4*/*55_=1.42, p=0.24), but differences in freezing damage between populations (F_4*/*59_=7.16, p*<*0.001). A post-hoc multiple comparison test (*α*=0.05, FDR corrected following (Benjamini & Hochberg, 1995)) combined the populations Juval, Laatsch and Vioz/Coro into a homogeneous subset with stronger leaf damage and the populations Terz and Finail in a subset with higher frost resistance (Figure 2). The differentiation suggests adaptation to local temperature regimes, but since both subsets include at least one high and one low altitude population and originated from different drainage systems, frost resistance in these populations is not strongly correlated with altitude nor with physico-geographic proximity.

**Figure 2:**
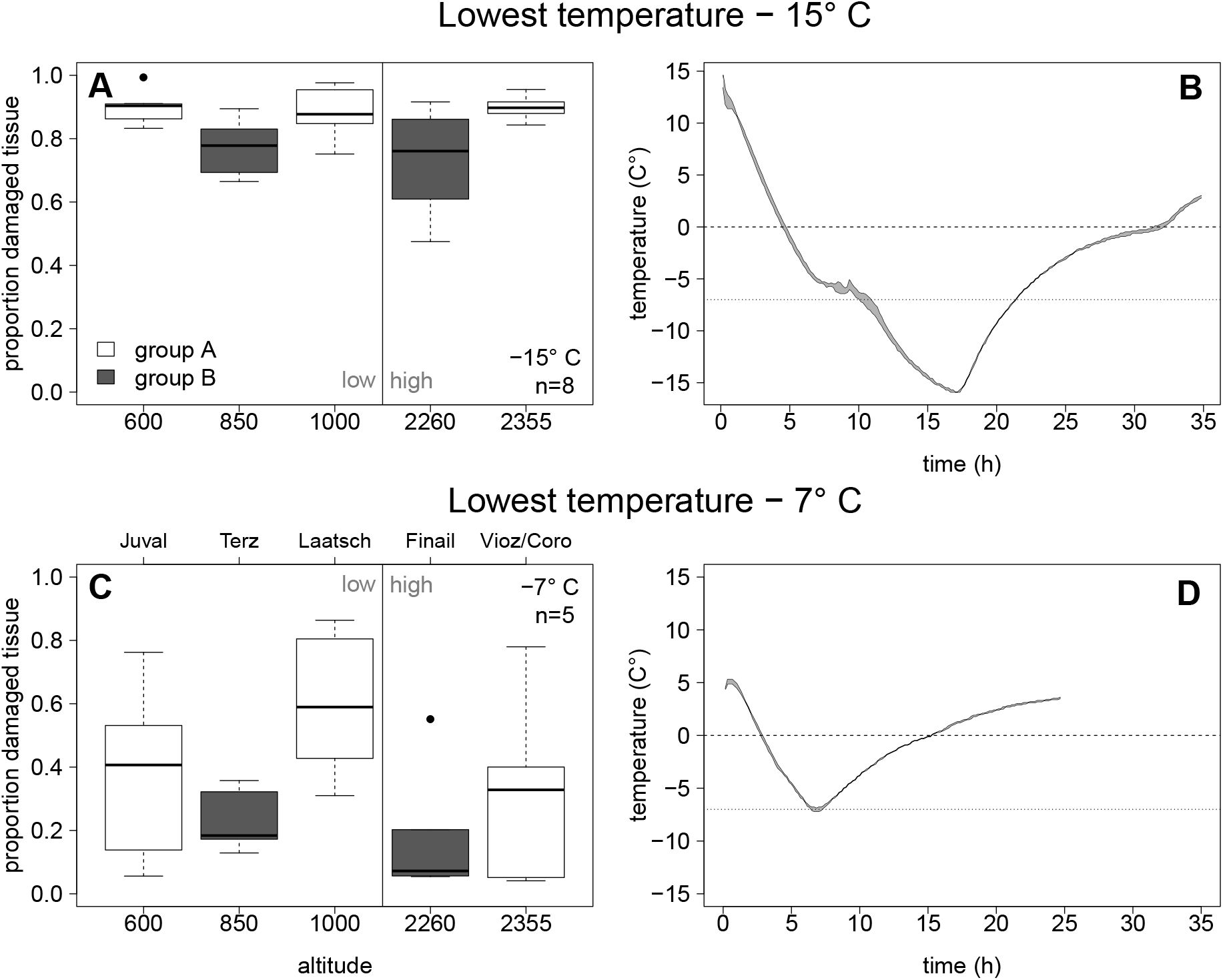
Freezing damage measured as cytosol leakage of destroyed tissue after freezing down to -15° C (A) and -7° C (C) as a fraction of cytosol leakage from completely destroyed tissue (*n* = 8). The vertical line divides the figure into the low altitude and the high altitude populations. Group A and B indicate homogeneous population subsets (*α* = 0.05) after correction for false discovery rate. (B) and (D) show the logged temperature as area between the measurements of two data loggers positioned above and below the probes.

As second phenotypic trait we investigated response to different types and strengths of light irradiation measured as red coloration of leaves, which is caused by anthocyanin accumulation and represents a typical stress reaction of plants (Park *et al*, 2007). The population × treatment (light condition) interaction was highly significant (F8*/*28=42, p*<*0.001) indicating differential population response to light stress. A post-hoc multiple comparison test (*α*=0.01, FDR corrected following Benjamini & Hochberg (1995)) on the GLS model indicated that plants from four of the five populations developed more red tissue under light stress than in the other light conditions (Figure 3A). Conversely, plants of the highest population (Vioz/Coro) were green in all treatments (Figure 3A, D) but developed more dead tissue under the high light treatment in comparison to the low light and UV-B treatments (Figure 3B, D, population × treatment interaction F_8*/*28_=14, p*<*0.001).

**Figure 3:**
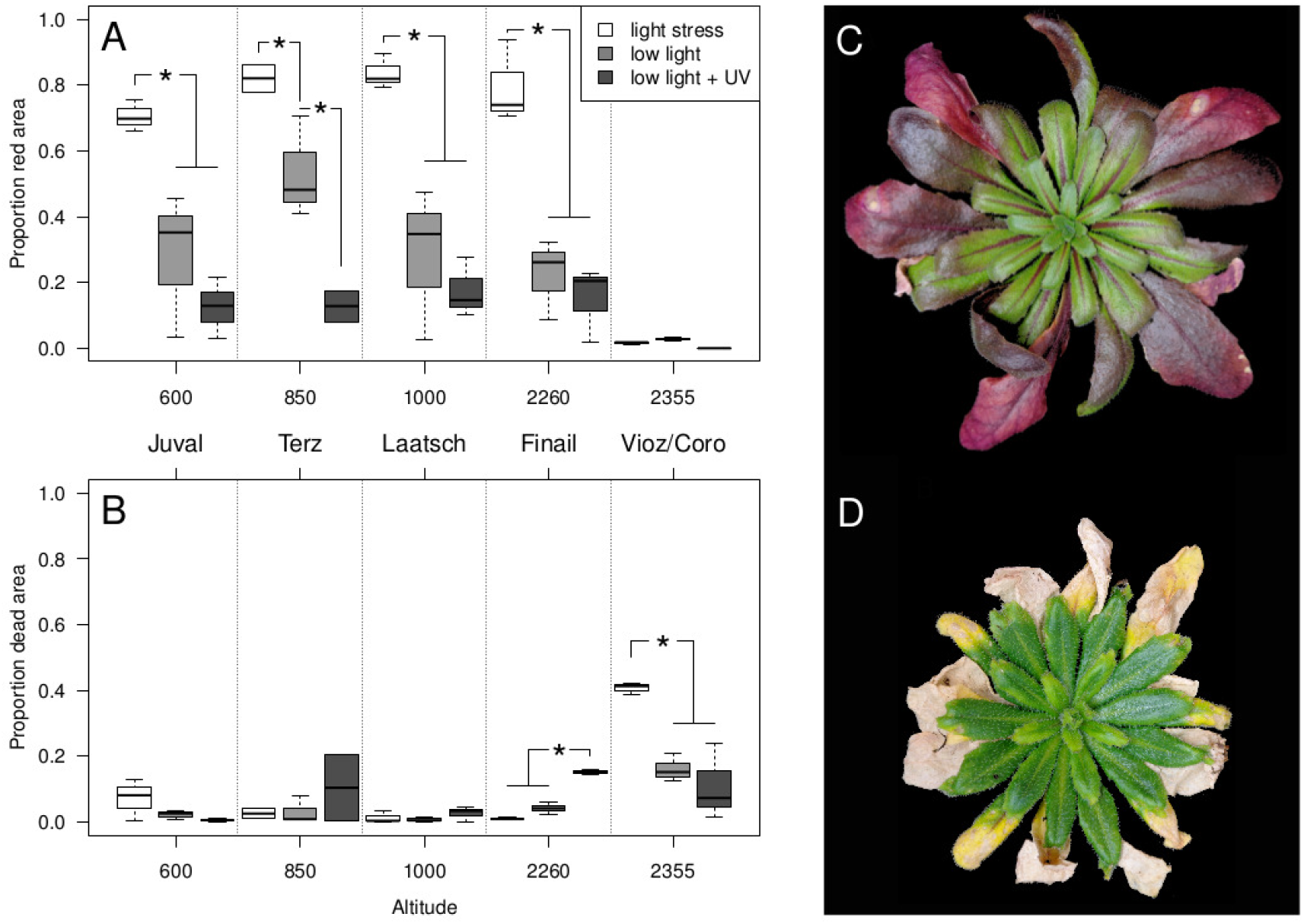
Phenotypic effects of 20 days of light stress treatment. A) Proportion of red tissue in light-stressed plants along the altitude gradient. B) Proportion of dead leaf tissue in light-stressed plants along the altitude gradient. Significant treatment differences at *p <* 0.001 or smaller (corrected for false discovery rate) are indicated with a star. C) A representative plant for the light stress treatment from the Finail population. D) A representative plant for the light stress treatment from the Vioz/Coro population.

An increased sensitivity to UV-B damage is expected in anthocyanin-deficient plants if the flavonoid pathway is affected simultaneously (Chalker-Scott, 1999). To test whether plants from high-altitude Vioz/Coro site showed an increased UV-B sensitivity, we exposed them to a gradient of increasing UV-B dosages over a period of three days and evaluated the proportion of dead tissue (Supplementary Methods). In contrast to the Finail population, which exhibited a superior UV-B resistance compared to the low altitude populations, plants from the Vioz/Coro population developed an increased proportion of dead tissue with higher UV-B dosage (binomial GLM; population × UV-B dosage F4*/*186=22, p*<*0.001; Figure S1, Table S1).

### Genetic diversity within and genetic relationship between populations

Pool sequencing of all five populations and subsequent mapping to the Col-0 genome resulted in a 19- to 50-fold median coverage of the nuclear chromosomes per population (Table 1). In each pool, about 95% of the reference genome sequence positions were sequenced at least once and a total of 3,075,350 SNPs were called. We first calculated nucleotide diversity, *π*, and Tajima’s *D* by accounting for pooling (Futschik & Schlötterer, 2010; Kofler *et al*, 2011a). As observed before (Clark *et al*, 2007; Cao *et al*, 2011), diversity was elevated in pericentromeric regions relative to chromosome arms (Figure 4A). The genome-wide average diversity of low altitude populations was high, while average Tajima’s *D* was close to 0 (Figure 4B, Table 1). In contrast, high altitude populations show a reduced nucleotide diversity and negative Tajima’s *D* values.

**Figure 4:**
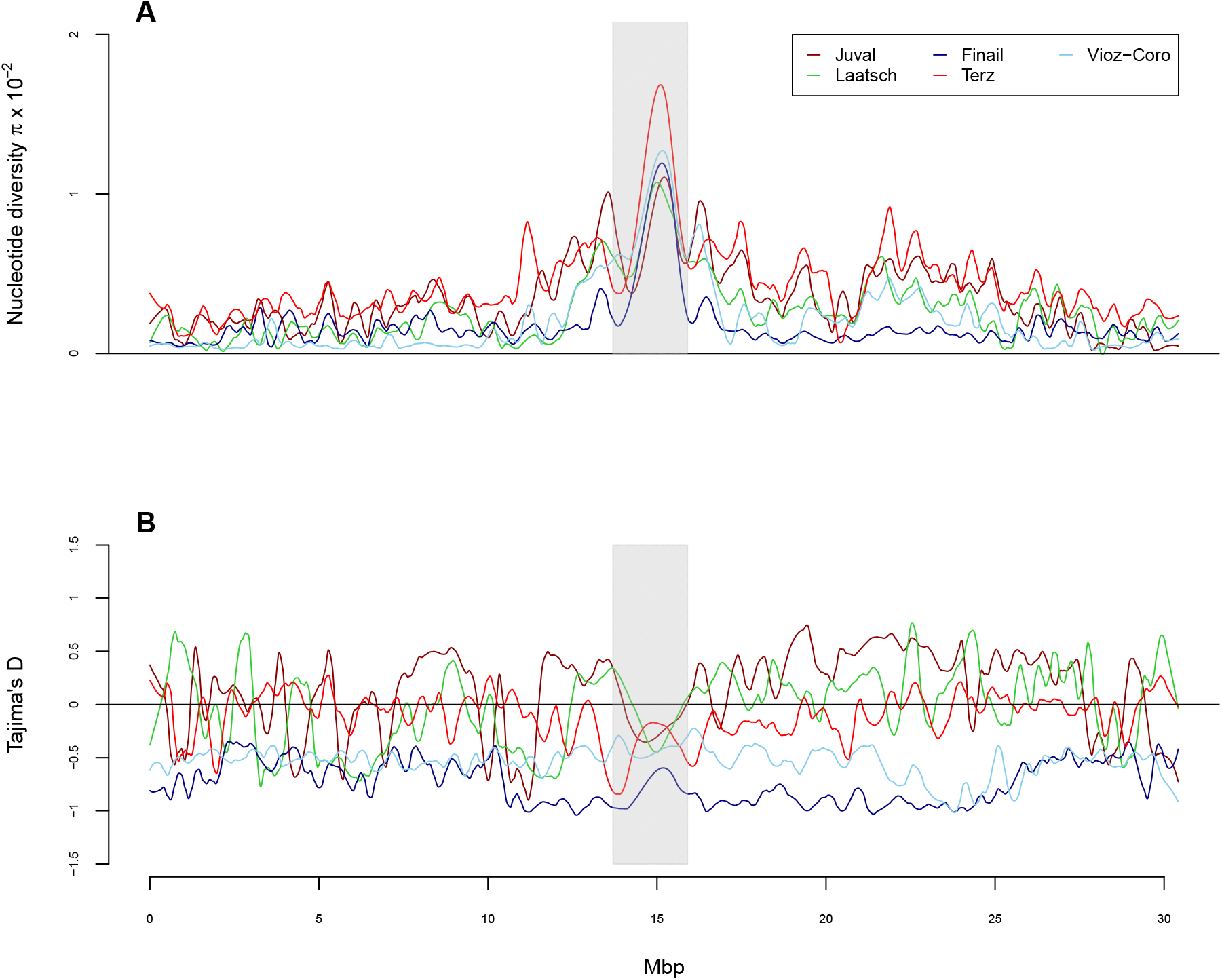
Sliding window analysis of (A) nucleotide diversity, *π* and (B) Tajima’s *D* for chromosome 1 in all five populations.

Allele frequency distributions were similar across populations with some interesting differences (Figure S4). All populations showed a large number of rare alleles and a substantial number of high-frequency or fixed derived alleles. The proportion of SNPs segregating at intermediate frequencies differed between populations (Table 1, Figure S4). SNPs with a strong putative functional effect (annotated as non-synonymous coding, stop lost, stop gained, start lost) segregate at much lower allele frequencies than other SNP types (Wilcoxon tests, *p <* 10^*-*7^ for all classes when compared to synonymous SNPs in each population). Allele frequencies of deleterious nsSNPs (as classified by MAPP) were lower than of tolerated nsSNPs in all populations (Wilcoxon tests, *p <* 10^*-*5^), indicating that purifying selection acted on this class of alleles. The ratio of nsSNPs to sSNPs, however, did not differ between high and low altitude populations (Fisher’s exact test, *p >* 0.05) suggesting that the strength of selection against deleterious polymorphisms was similar in high and low altitude populations.

The pairwise correlations of allele frequencies derived from the covariance matrix estimated by Bayenv2.0 and pairwise *F*_*ST*_ values (Table S2) revealed that both matrices were correlated (Mantel-test, *r* = *-*0.63, *p* = 0.035). Pairwise *F*_*ST*_ values were similar between all pairs of populations, although the maximal distance was observed between Vioz/Coro and Terz populations, which belong to the same drainage system. This pair also showed the lowest correlation of allele frequencies as estimated by Bayenv2.0. Generally, the correlation estimates gave a higher variation than the *F*_*ST*_ values across population pairs (Table S2), because Bayenv2.0 accounts for different sequence coverage in the correlation matrix.

Neighbour-Joining trees generated from the two distance matrices indicate a closer relationship between the two high altitude populations than to the low altitude population from the same drainage system (Figure 5). The *F*_*ST*_-based tree is almost star-like with short internal branches, whereas the tree derived from the covariance matrix shows longer internal branches and a stronger clustering of population pairs. We observed essentially the same clustering with a TreeMix analysis based on 1.07 million SNPs with a minimum coverage of 20 per population (Figure 5C). Bootstrapping results of the TreeMix analysis show a 77% confidence for the Juval and Terz population group and a migration event with 73% confidence from the Finail to the Juval population. The four-population test (Reich *et al*, 2009) rejected all possible tree topologies (all |*z*| *>* 3, which corresponds to *p <* 0.001), but the lowest |*z*| was observed for the tree that groups the two high altitude populations together (Table 2). The positive test statistic for this tree suggests some admixture in at least one drainage system, most likely between Finail and Juval, consistent with the TreeMix results.

**Figure 5:**
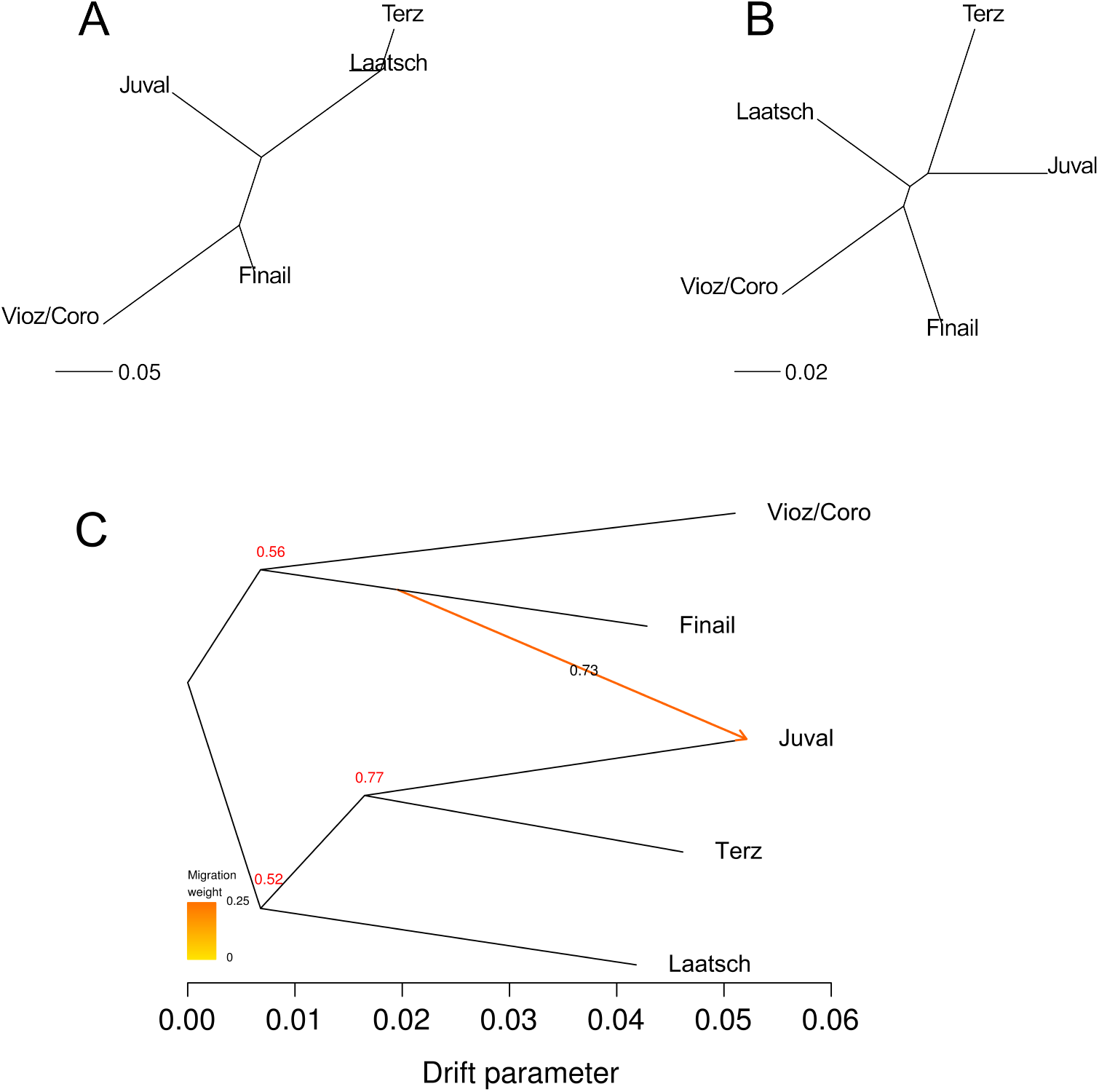
Relationship between populations based on measures of pairwise genetic differentiation and on allele frequencies. Branch lengths reflect the extent of genetic differentiation. (A) NJ tree of the correlation matrix obtained with Bayenv2.0. (B) NJ tree of the matrix of pairwise *F*_*ST*_ values. (C) Tree inferred with TreeMix of the five alpine populations with migration events based on 1.07 Million SNPs with a coverage of *>* 20. The migration event is coloured according to its weight which represents the percentage of alleles originating from the source. Numbers at branching points and on the migration arrow represent bootstrapping results based on 1,000 runs.

**Table 2:**
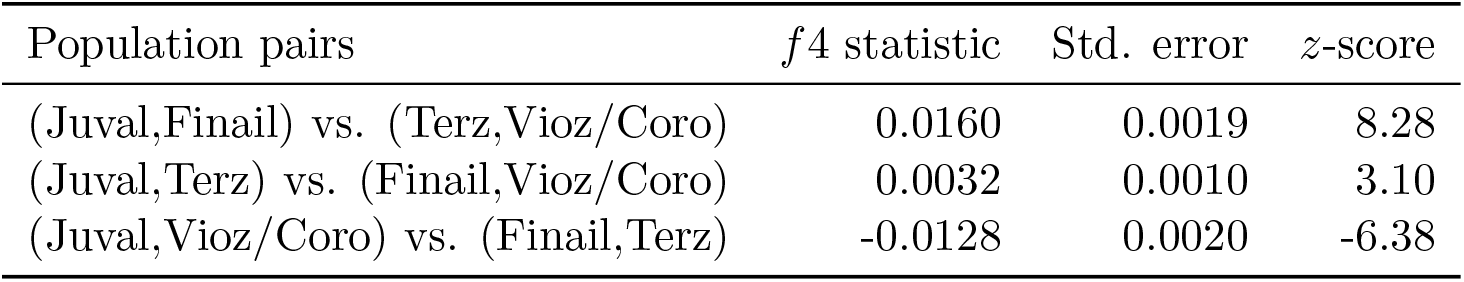
Four-population test of two high and two low-altitude populations. It tests the probability that the relationship of four populations can be explained by one of three possible tree topologies without migration. The test was conducted with 1.95 Million SNPs with a minimum coverage of 20 in each population.

| Population pairs | $f_4$ statistic | Std. error | $z$-score |
| --- | --- | --- | --- |
| (Juval,Finail) vs. (Terz,Vioz/Coro) | 0.0160 | 0.0019 | 8.28 |
| (Juval,Terz) vs. (Finail,Vioz/Coro) | 0.0032 | 0.0010 | 3.10 |
| (Juval,Vioz/Coro) vs. (Finail,Terz) | -0.0128 | 0.0020 | -6.38 |

### Modelling the demographic history with ABC

The tree-based analyses suggested a common origin of the two high altitude populations. We therefore considered each pair of low and high altitude populations from the same drainage system as an independent replicate of the same demographic process to estimate divergence time between high and low altitude populations. In both analyses the exponential growth model with bidirectional migration gave posterior probabilities of almost 100% (Table S4) with very similar parameter estimates. The effective population size of the high altitude populations was smaller than of the low altitude populations after the population split (Table 3), with up to a three-fold difference in population size. The 90% HPDI of the split time in the models ranged from 14,800 to 260,000 years before present, and the interval was almost identical between the two population pairs (Table 3).

**Table 3:** Posterior probability of the parameters of the best-fitting demographic model (exponential growth, with migration). The probability for each parameter was inferred by ABC.

| Parameter | Juval - Finail |  | Terz - Vioz/Coro |  |
| --- | --- | --- | --- | --- |
|  | Mode | 90% HPDI | Mode | 90% HPDI |
|  | *Exponential growth, with migration* |  | *Exponential growth, with migration* |  |
| $N_{\text{low alt.}}$ | 195,457 | 168,005 - 199,983 | 196,573 | 176,779 - 199,988 |
| $N_{\text{high alt.}}$ | 71,221 | 19,460 - 166,695 | 77,039 | 18,275 - 169,580 |
| $T_{\text{split}}$ | 70,650 | 18,160 - 261,636 | 77,491 | 14,892 - 260,579 |

### Highly differentiated SNPs between high and low altitude populations

To identify candidate loci for altitude specific adaptation, we correlated allele frequencies of populations with altitude using Bayenv2.0. The *Z* (Figure S5) statistic (Günther & Coop, 2013) was calculated for 400,231 SNPs in total after quality control (see methods section). A total of 25 SNPs showed the highest score possible for *Z* corresponding to a strong support for a non-zero correlation (i.e. *Z* = 0.5; Table S6). One non-synonymous SNP among them was found in the gene AT5G17860, which codes for the Calcium exchanger 7 (CAX7) protein located on chromosome 5 (Figure 6A, B). A region about 5 kb upstream of this gene showed no sequence coverage in both high altitude populations and the Laatsch population (Figure 6B, C). This pattern indicates that the Col-0 reference haplotype is not present in these populations, either because of a deletion or the presence of a highly divergent haplotype of a length of about 25 kb. The whole region was flanked by SNPs with high *Z* values and strong differentiation between high and low altitudes (Figure 6A). To test whether this pattern resulted from insufficient local sequence coverage, we analysed all 33,268 TAIR10 genes with sequence data in at least one population. Only 15 genes showed a similar pattern, which we defined as a per base coverage below the 5th percentile in high altitude and above the 5th percentile in low altitude populations. This low probability to find similar presence-absence variation in our sequence data supports a deletion in the high altitude populations. Of these 15 genes, 12 genes were annotated as unknown protein, protein of unknown function, pseudogene or transposon; one gene encodes a ribosomal protein, one gene a cytochrome P450 family protein and the last one is an NBS-LRR receptor (AT5G48770).

**Figure 6:.**
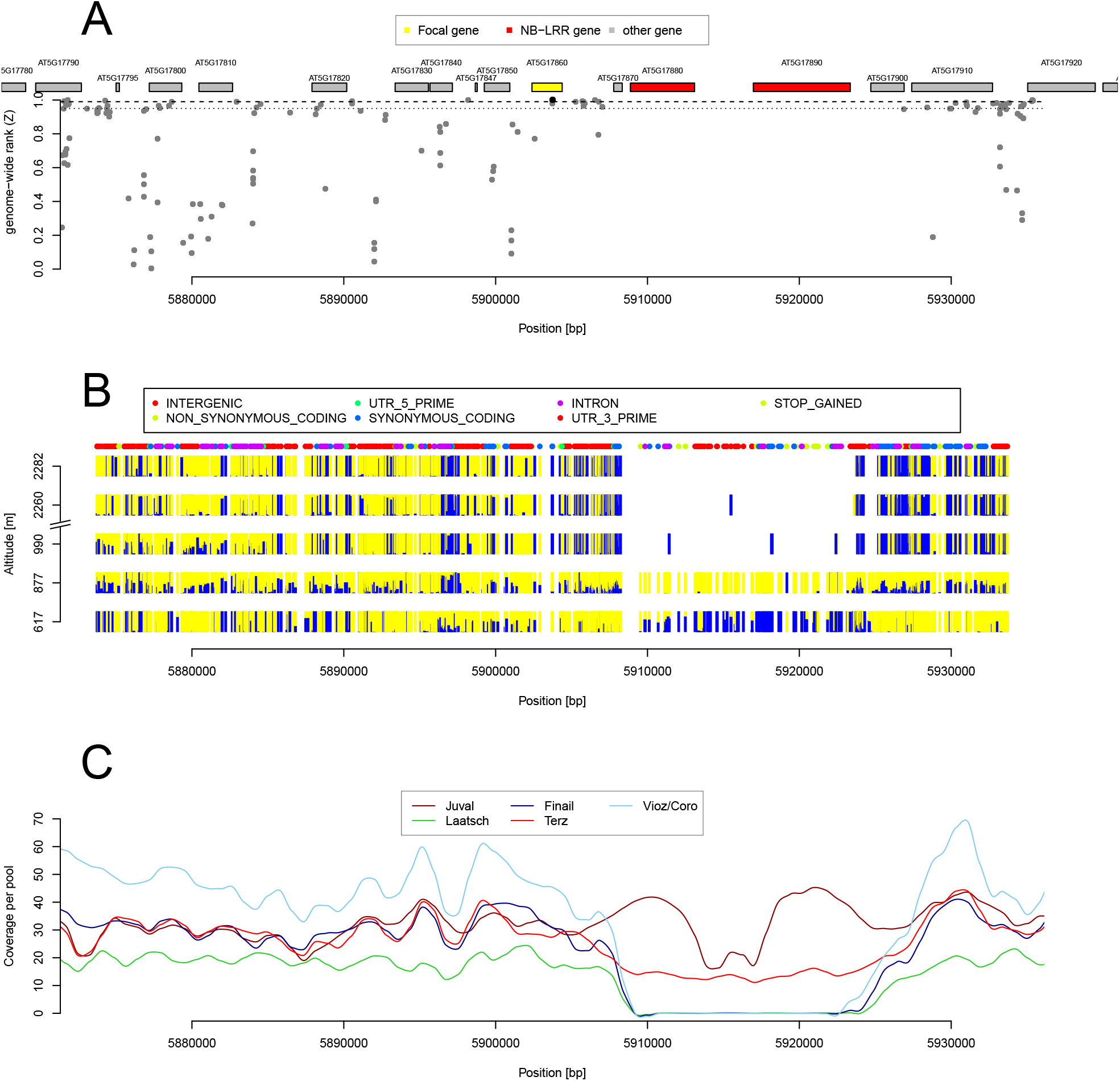
Example of a candidate gene (AT5G17860, encoding for *CAX7*) in which allele frequences were highly differentiated between high and low altitude populations. (A) Plot of the genome-wide relative rank of the *Z* statistics. High values indicate a strong differentiation. (B) Plot of the relative allele frequencies of the two alleles segregating at a polymorphic site. The height of the bar is proportional to the allele frequency. (C) Sequence coverage of the genomic region in the five different populations.

A second candidate gene encodes the Eps15 homology domain protein AtEHD1 (AT3G20290) where a single SNP located in an intron shows a high Z-score (*Z* = 0.5) but is flanked in a region of at least 50 kb by additional SNPs highly associated with altitudes. This region includes two additional genes downstream of the *AtEHD1* gene (Figure S7). In general the proportion of fixed alleles in the high-altitude populations is much larger than in the low-altitude populations. Here, only a single SNP shows the highest correlation with altitude, however, across a large region the two high-altitude populations were strongly differentiated from the other two populations.

A third example of an outlier region is located on chromosome 3 around a highly differentiated SNP in the gene AT3G07130 which encodes the purple acid phosphatase 15 (Figure S8). In this region, only the Vioz/Coro population at the highest altitude has a large number of fixed alleles and is strongly differentiated from the other four populations, which is consistent with a footprint of selective fixation of alleles in this population.

### Enrichment analysis of highly differentiated genes

To test whether functional groups of genes show a higher level of differentiation, we considered the top 1% of SNPs as highly differentiated SNPs and conducted an enrichment analysis with Gowinda. This analysis did not uncover enriched gene sets after correction for multiple testing with the exception of gene sets obtained from a literature collection (Lai *et al*, 2012) (Table S11). Nevertheless, categories with nominal p-values ≤ 0.05 included annotations expected to be involved in adaptation to different altitudes (Tables S7 and S8). For example, biological processes like response to light stimulus, heat acclimation, defence signalling pathway, flavonol biosynthetic process and biochemical processes like superoxide radical degradation, glucosinolate biosynthesis, flavone and flavonol biosynthesis and anthocyanin biosynthesis are some of the processes which are likely involved in adaptation to altitude. We also annotated the functional effects of highly differentiated SNPs (Figure S6). Genes highly differentiated between high and low populations were enriched for genic SNPs because most SNPs were located in introns and UTRs, but not in coding regions. If genic but non-coding SNPs were involved in adaptive differentiation, selection should have preferentially acted on regulatory processes rather than on protein function. Such a hypothesis was supported by an enrichment analysis of highly differentiated SNPs in known predicted transcription factor binding (TFB) sites. According to the AGRIS database (Yilmaz *et al*, 2011) 3.3% of all SNPs were located in TFB sites, but highly differentiated SNPs were enriched in TFB sites although effect sizes were small (Bayenv2.0 candidates: 3.75%; Fisher’s exact test, *p* = 0.03711).

## Discussion

### Presence of phenotypic variation in low and high altitude populations

We identified phenotypic variation in two ecologically relevant traits in the five populations but their spatial distribution did not show a strong correlation with altitude. Due to the large difference in altitude between populations (*>*1,000 m) and a decrease of average temperature by 0.6 °C per 100 m altitude (Barry, 1992) we expected strong selection for freezing tolerance at high altitude. Although we noted a significant variation in freezing tolerance, populations with similar frost resistance levels did not originate from the same altitude or water drainage (river) system. A possible explanation is that selection for cold tolerance is mainly influenced by microclimatic conditions which may be strongly influenced by local topology. The high altitude population Vioz/Coro is found on South-exposed rocky outcrops at protected sites underneath rock overhangs. Temperature logger data from the two high altitude sites suggest that the Vioz/Coro site is less prone to freezing during winter (Figure SS2). In addition, high alpine *A. thaliana* populations may switch to a summer annual life habit allowing the plants to avoid the period with high risk of freezing damage (Luo *et al*, 2014). Our observations of plantlets at the different localities suggest that the Vioz/Coro population consists of summer annuals, while at the other four sites plantlets germinate in the fall and persist throughout the winter.

The spatial pattern of response to light and UV-B stress differed from frost tolerance. The population at the highest altitude (Vioz/Coro) showed a lack of anthocyanin accumulation under light stress. Anthocyanins protect against cell damage by light-induced photooxidation (Chalker-Scott, 1999; Powles, 1984) and the plants from the Vioz/Coro population exhibited tissue bleaching in old roset leafs under light stress and were more sensitive to extended elevated UV-B levels. Flavonoid-deficient mutants of *A. thaliana* show a similar phenotype (Li *et al*, 1993). Solar radiation at open sky and fluctuations in radiation intensity increase with higher altitude (Blumthaler *et al*, 1997; Körner, 2007) and high altitude populations need to adapt to more intense UV-B and light conditions. For example, the Shahkdara accession of *A. thaliana* was collected at 3,063 m a.s.l. and exhibited a constitutive UV-B protection (Biswas & Jansen, 2012). We therefore suggest that the deficiency in anthocyanin accumulation in the Vioz/Coro population is not an adaptation to high altitudes but a loss-of-function phenotype resulting from genetic drift at the range margins (Eckert *et al*, 2008).

In summary, the spatial patterns of variation in the two traits do not support a simple pattern of adaptation to altitude because we expected both high altitude populations to show a similar phenotype and the demographic analysis suggested that there was sufficient time available for adaptation to high altitude. Possible explanations are that a difference of 1,700 m altitude between sampling locations does not represent a strong selection gradient or that the microenvironment or random drift and metapopulation dynamics of small demes confounds clinal trait variation.

### Patterns of genetic diversity and population differentiation

In *A. thaliana*, levels of genetic diversity differ strongly among populations at small and large geographical scales (Bomblies *et al*, 2010; Cao *et al*, 2011). A low genetic diversity limits the potential to respond to selection and may lead to the accumulation of deleterious alleles (Alleaume-Benharira *et al*, 2006). In our sample, both high altitude populations exhibited reduced genetic diversity and genome-wide negative Tajima’s *D* values that indicate a deviation from a standard neutral model. This pattern may result from bottlenecks with subsequent population growth (Simonsen *et al*, 1995) as expected after the colonization of new habitats like high altitude sites at the outer vertical limit of a species distribution. However, we did not observe an increase of deleterious amino acid polymorphisms in high altitude populations that would accompany such a bottleneck due to a reduction of purifying selection (Günther & Schmid, 2010).

Although diversity estimates differed significantly among the five populations, we found little evidence for a strong population structure. The average pairwise *F*_*ST*_ value of 0.128 was three times smaller than in local populations near Tuübingen, Germany (*F*_*ST*_ = 0.52; Bomblies *et al*, 2010) and smaller than the smallest observed pairwise *F*_*ST*_ in a sample of Swiss alpine *A. thaliana* populations that were based on 24 SSR loci (Luo *et al*, 2014). In addition, pairwise *F*_*ST*_ values were very similar for all pairs of populations, consistent with a low number of unique SNPs per accession and a high level of LD in a different sample of South Tyrolian accessions (Cao *et al*, 2011). The demographic history of numerous alpine plant species was influenced by glaciation which caused repeated cycles of retreats into refugia and colonization of newly available habitats in interglacial periods (Schönswetter *et al*, 2005). Refugial populations in alpine valleys or local refugia on mountain tops (nunataks) above glaciers may have contributed to a strong genetic differentiation between subpopulations. A low differentiation and a star-like relationship with gene flow between populations (e.g., the low-altitude Juval and the high-altitude Finail populations; Figure 5) of our five populations is not consistent with such a scenario and instead suggest that these sites were colonized recently from refugia at the edge or outside the alps. On the other hand, the two high altitude populations were more closely related to each other than to the respectice low-altitude populations from the same river drainage. This may reflect an origin from the same (mountaintop) refugium or a recent dispersal by animals. Generally, a rapid postglacial recolonization of alpine habitats by *A. thaliana* is possible because of the mainly self-fertilizing mode of reproduction and dispersal by animals like mountain goats. The high altitude populations were found only at highly disturbed resting sites of mountain goats.

To investigate the age of the split of low and high-altitude populations, we estimated demographic parameters for different demographic models with ABC (Beaumont *et al*, 2002). Both pairs of low and high altitude population supported a model with a smaller effective population size in high altitude populations caused by a bottleneck and subsequent population growth. The 90% HDPI intervals for population split times ranged between 14,000 and 260,000 years before present. This includes time before and after the last glacial maximum (LGM) as deglaciation on the Northern Hemisphere began about 19-20 thousand years ago (Clark *et al*, 2009). Consequently, our ABC analysis did not allow to answer the question whether the split between low- and high-altitude populations occurred before or after the LGM, though the probability is higher that it occurred before. The wide HDPI 90 interval may result from a limitation of pool sequencing because summary statistics used for ABC were limited to measures of gene frequencies and did not include haplotype-based statistics, which provide additional power for inferring population split times and admixture levels by quantifying the length of shared fragments (Sved *et al*, 2008; Sankararaman *et al*, 2012; Hellenthal *et al*, 2014).

### Patterns of genetic variation and local selective sweeps

In *A.thaliana* only few species-wide selective sweeps were found so far (Clark *et al*, 2007; Childs *et al*, 2010; Cao *et al*, 2011; Horton *et al*, 2012), and the overall frequency of adaptive evolution is very low or absent based on genome-wide polymorphism data (Gossmann *et al*, 2010; Slotte *et al*, 2011) although evidence for local adaptation was found in several studies (Fournier-Level *et al*, 2011; Hancock *et al*, 2011; Agren & Schemske, 2012; Méndez-Vigo *et al*, 2011; Long *et al*, 2013; Huber *et al*, 2014). Therefore, searching for signatures of selection at smaller geographic scales along environmental gradients may identify phenotypic and genomic footprints of adaptive evolution. We identified candidate genes for high-altitude adaptation with an outlier approach. It should be noted that this is no formal test of selection and candidate genes require further validation (Pavlidis *et al*, 2012).

Patterns of genetic variation at some candidate genes suggested that targets of selection were difficult to elucidate because large genomic regions (*>*50k) around significantly associated SNPs involving multiple adjacent genes showed a population-specific sweep pattern (Figures 6, S7 and S8). The comparison of the three genomic regions showed that despite an overall low level of genetic diversity in the alpine populations, complex patterns of genetic diversity that involved presence/absence variants (PAVs) and regions of high nucleotide diversity were observed. For example, in our sample, a single SNP was significantly differentiated between low and high altitude populations and some neighboring SNPs also showed a clinal pattern (Figures 6). It was previously identified in *Arabidopsis lyrata* as a candidate selection target for adaptation to soils with a low Ca:Mg-ratio (Turner *et al*, 2008) and *A. thaliana* is generally found on siliceous rock with low Ca content. However, in the direct neighbourhood, although pool sequencing did not allow the analysis of haplotypes, patterns of sequence coverage strongly suggested that a PAV comprising two well characterized NBS-LRR genes, *CSA1* and *CHS3*, may have contributed to local adaptation since the deletion was fixed in the high altitude and Laatsch populations, whereas it segregated in the low-altitude populations (Figure 6). In addition both genes showed a PAV across the whole species because *CSA1* was not covered by sequence reads in 11 and *CHS3* in 9 out of 18 high coverage reference genomes of a diverse set of accessions (Gan *et al*, 2011). PAVs are abundant across the genome of *A.thaliana* (Tan *et al*, 2012), particularly in resistance genes that include NBS-LRR genes (Bergelson *et al*, 2001; Shen *et al*, 2006; Guo *et al*, 2011; Karasov *et al*, 2014). It remains open whether selection or genetic drift contributed to the fixation at these genes. They are involved in pathogen response, but *CSA1* additionally plays a role in shade avoidance and response to red light (Faigón-Soverna *et al*, 2006) and knockouts of *CHS3* cause chilling-sensitive plants (Schneider *et al*, 1995). A comparison of sequence coverage pattern against all genes annotated in TAIR10 showed, however, that a similar deletion pattern was observed in only 15 genes that included other NBS-LRR genes or genes of unknown function.

The pattern of diversity at the *AtEHD1* gene (AT3G20290) may represent a population-specific selective sweep because of numerous SNPs with a high *Z*-score in this region. The same alleles are fixed in the two high-altitude populations and the diversity is strongly reduced over a range of about 50 kb (Figure S7). In contrast, the low altitude populations appeared to be highly polymorphic in this region. The identification of putative selection targets requires further study, but it is noteworthy that *AtEHD1* increases tolerance to salt stress (Bar *et al*, 2013).

We observed another pattern of genetic diversity in the gene encoding the purple acid phosphatase 15 (AT3G07130) where only the highest population was differentiated (Figure S8). This gene may affect phosphorus efficiency by remobilizing inorganic phosphate from organic phosphate sources (Wang *et al*, 2009) and is a plausible selection target because phosphorus was highest in the highest population (Table S5). Taken together, diversity patterns in several genomic regions suggest that adaptation to soil conditions may have played a role in the history of these populations (Baxter *et al*, 2010).

Changes in light conditions and temperature differences are among the expected environmental differences between altitudes. We did not identify genes known to be involved in frost resistance in the GO enrichment analysis (Table S7), which corresponds to the patterns of phenotypic differentiation. However, genes associated with heat acclimation were among the identified candidates, which may relate to the observation that high temperatures in late spring terminate the reproductive cycle in all low altitude populations by causing plant death. In contrast, mild temperatures at high altitudes allow reproduction throughout the summer suggesting selection for heat acclimation may be relaxed (personal observation: C. Lampei). Furthermore, in several genes involved in shade avoidance and response to light stimulus as well as mutations in the anthocyanin (Table S8), flavonol and phenylpropanoid biosynthesis pathways (Table S7) were correlated with altitude. This pattern is consistent with phenotypic differentiation, although the anthocyanin deficiency may constitute a loss of function mutation in the highest population, indicating that genetic differentiation at some genes may not result from past selection.

In a recent genome-wide scan in *A. halleri*, genes associated with red or far red light response were differentiated with altitude in two independent altitude gradients in Japan (Kubota *et al*, 2015). Further, genes involved in defence response were also enriched in an Alpine altitude gradient study in the same plant (Fischer *et al*, 2013). Notably, the enrichment analysis did not identify any soil related categories among the nominally significant annotations, which is in strong contrast to the annotation of the top 25 highly differentiated genes. This corresponds to the observation by Fischer *et al* (2013) that the GO enrichment analysis did not match with outlier correlations with environmental variables.

### Methods to identify selection along altitudinal gradients

Pool sequencing is a cost-efficient method to characterize genetic variation using allele frequency estimates. Reliable estimates of population-specific allele frequencies require a sufficient sample size and sequence coverage (Schlötterer *et al*, 2014). Our sequenced pools fulfill recommendations for pool sequencing and we expect that the observed patterns of genetic diversity resemble genomic differences and not artefacts of the method. Both our phenotypic and genomic analysis identified substantial variation along altitudinal gradients. Studies of local adaptation need to consider the effect of sampling design on the probability to identify false positives (Meirmans, 2015). Simulations suggest that studies of local adaptation should be based on several pairs of populations that differ in the environmental variable of interest (Lotterhos & Whitlock, 2015). We included two population pairs to have to samples of putative adaptive processes and demographic history. The closer relationship of the high-altitude populations, however, suggest that genetic variation shares a joint history and are not entirely independent. For this reason, the power to identify targets of altitudinal adaptation increases with more population pairs, if local populations are small and isolated and therefore prone to genetic drift. Future studies of altitudinal adaptation in *A. thaliana* should therefore be based on a denser sampling of populations and include additional phenotypic traits (Montesinos-Navarro *et al*, 2011; Luo *et al*, 2014). Given the patchy and highly disjunct distribution of *A. thaliana* in the high Alps, this might require systematic sampling over a larger geographic area to disentangle demography from natural selection (Meirmans *et al*, 2011).

## Conclusion

The phenotypic variation in response to two environmental variables, which predictably change with altitude, was not consistent with selection for high-altitude adaptation in our *A. thaliana* populations from the North Italian Alps. Microevolutionary responses to local topology at the population sites, founder effects and small population sizes at high altitude may overrule mean altitudinal change in environmental variables. In contrast, several genomic regions showed patterns of genetic variation consistent with population-specific selection-driven sweeps, that were associated with observed environmental differences (e.g. soil parameters). Furthermore, the detection of outlier genes suggested some plausible targets positive selection. These candidate genes can be tested at a functional level in future studies to test their potential role in adaptation to high altitude. Patterns of genetic diversity indicated a closer relationship of the high altitude populations that may reflect a common demographic history during glacial-interglacial cycles.

## Acknowledgements

We thank Elisabeth Kokai-Kota for lab assistance, Hinrich Bremer for access to a conductivity meter, Dr. Nikolaus Merkt for access to a UV-light meter and Fabian Freund for statistical discussions. This work was funded by the ESF EUROCORES project EpiCol (SCHM 1354/4-1), the German Federal Ministry for Education and Research (BMBF) within the AgroClustEr Syn-breed—Synergistic plant and animal breeding (FKZ0315528D), and BMBF Plant2030 project RYE-SELECT (FKZ 0315946E).

## Data availability

The raw sequence data were deposited in the Short Read Archive (XXXXX). SNP calls and other downstream sequence analyse data were deposited with with all phenotypic measurements and the R and Python scripts used for the analysis at DataDryad under accession number XXXXX.

## Supplementary Methods

### Resistance to high UV-B radiation

To test resistance to higher UV-B dosages we subjected 44 days old plants (10 pots and genotypes per population) which were still small due to 19 days of vernalization (4C and short day (8h light)) to increasing UV-B dosages of continuous UV-B light over 3 days. The UV-B levels were 0, 2.47, 6.39 and 15.16 *kJ m*^*-*2^ *day*^*-*1^. After this treatment plants were left to recover for 8 days to improve the separation of dead and alive biomass. All plants were evaluated manually with their identity hidden from the investigator. Dead and living tissue was separated for each pot and oven dried for 12 h at 90 C before weighting. A GLM with dead biomass odds as response variable and the fixed effects population and UV-B dosage was fitted with a quasibinomial error distribution (logit link) in R (DevelopmentCoreTeam, 2014, package:stats).

### Soil properties

In all sites soil samples of the top soil were collected. The soil samples were composed from about 40 individual samples per population site which were collected with a hand shovel from the uppermost 5 cm well spread across the distribution of the population. The soil was mixed, homogenized and analysed for standard parameters (Table S5).

### Supplementary Figures

**Figure S1:**
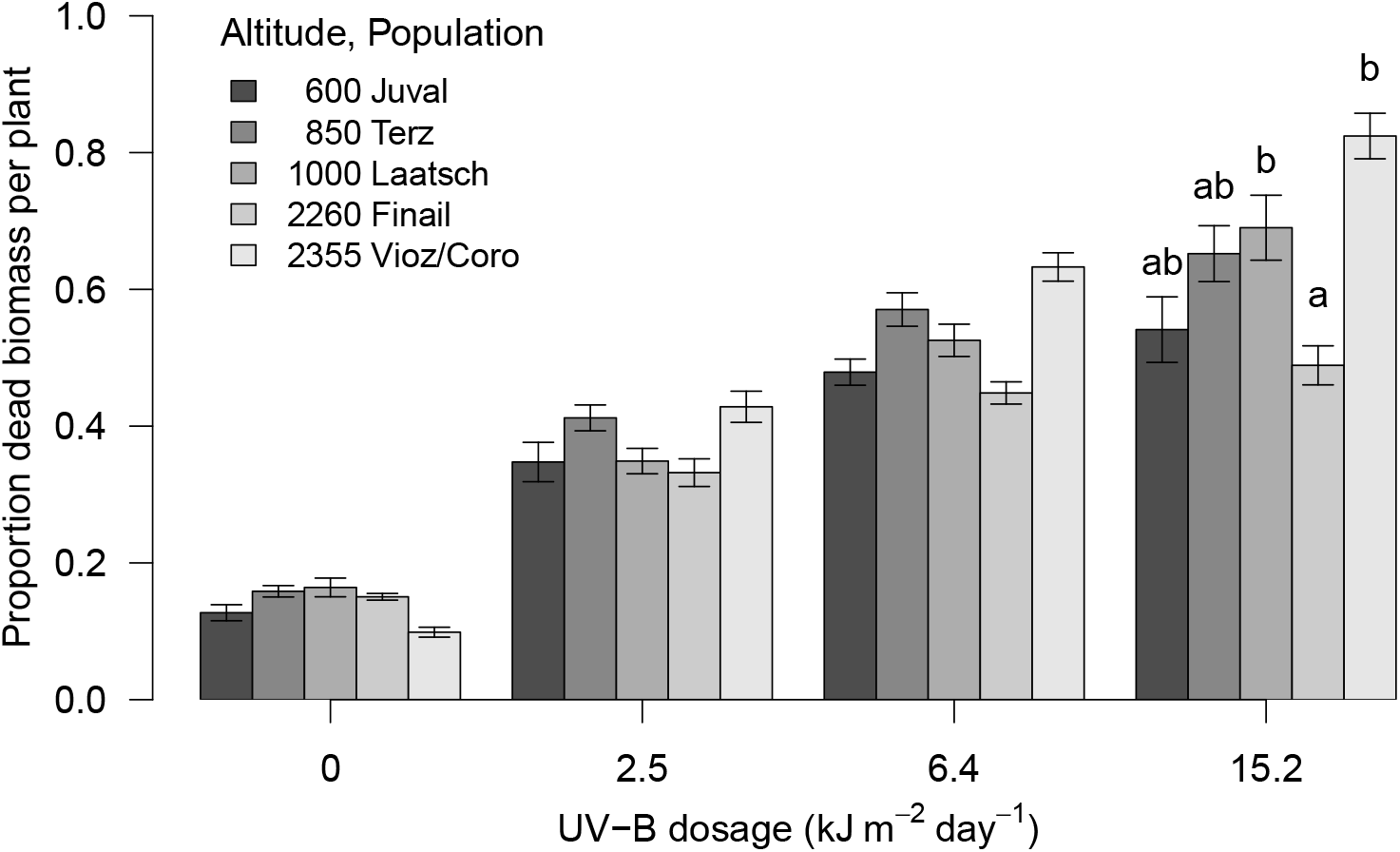
Proportion of dead tissue after 3 days continuous UV-B treatment with different dosages. Letters above error bars indicate homogeneous subsets after correction for false discovery rate for the highest UV-B dosage

**Figure S2:**
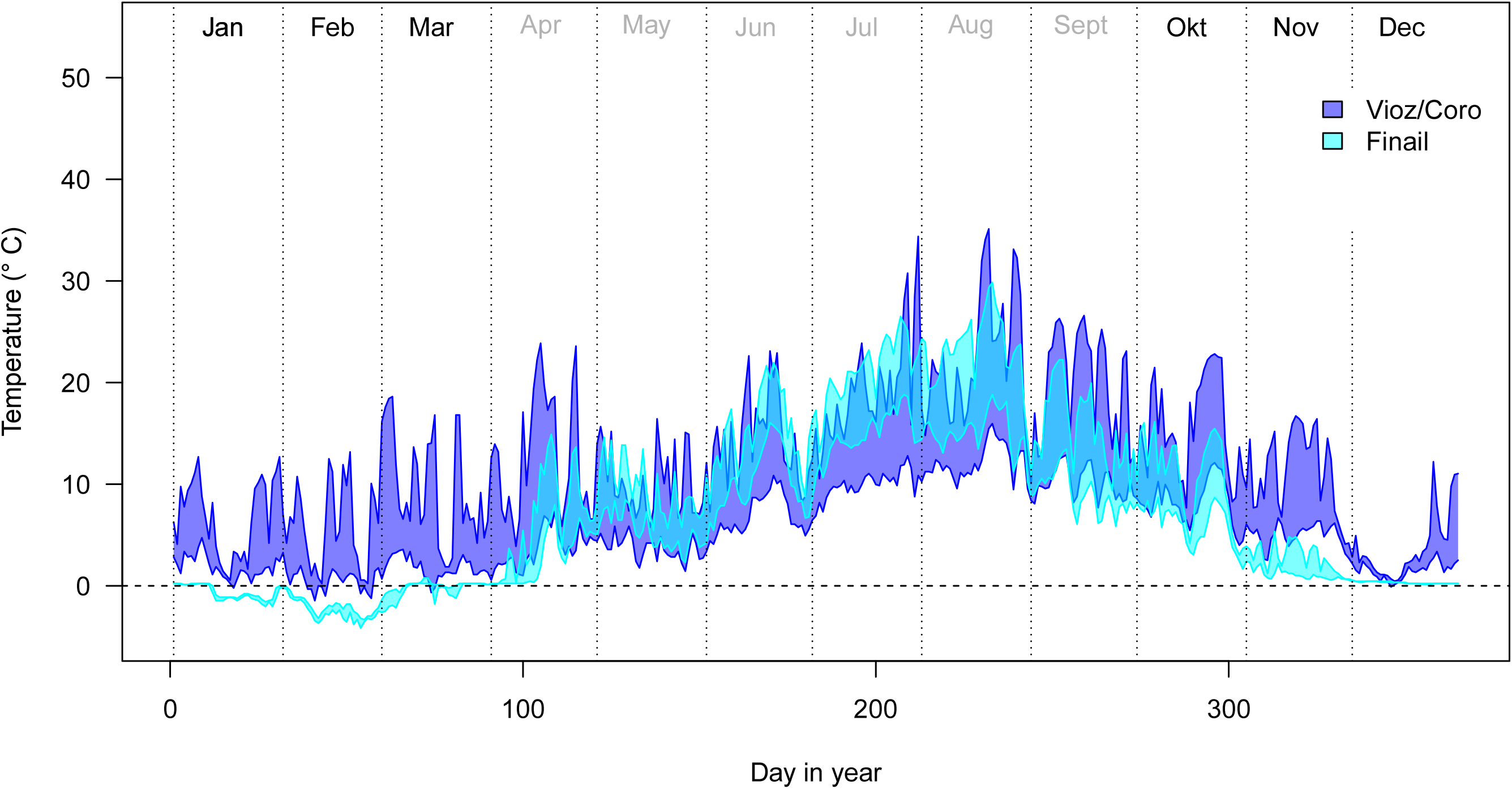
Daily minimum and maximum temperature for the top soil layer (< 5 cm) in the two high altitude sites

**Figure S3:**
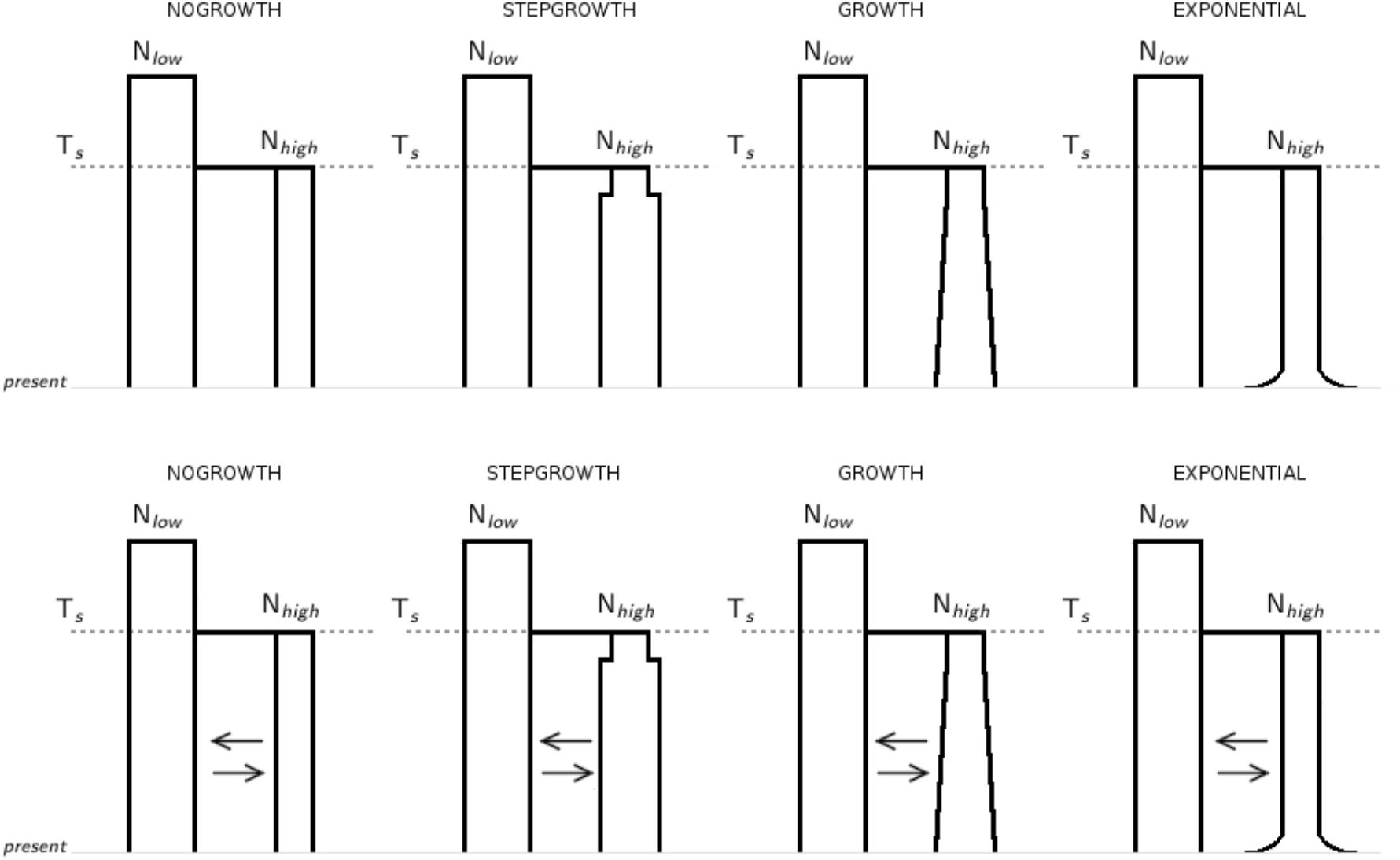
Four demographic models without and with migration investigated in this study. NOGROWTH - Model without the growth of the high altitude population after the split. STEPGROWTH - Model with one instantaneous size change of the high altitude population after the split. GROWTH - Model with simulated growth of the high altitude population after the split. EXPONENTIAL - Model with an exponential growth of the high altitude population sometime after the split. T_*s*_ - split time. N_*low*_ and N_*high*_ - effective population size of the low and high altitude population, respectively.

**Figure S4:**
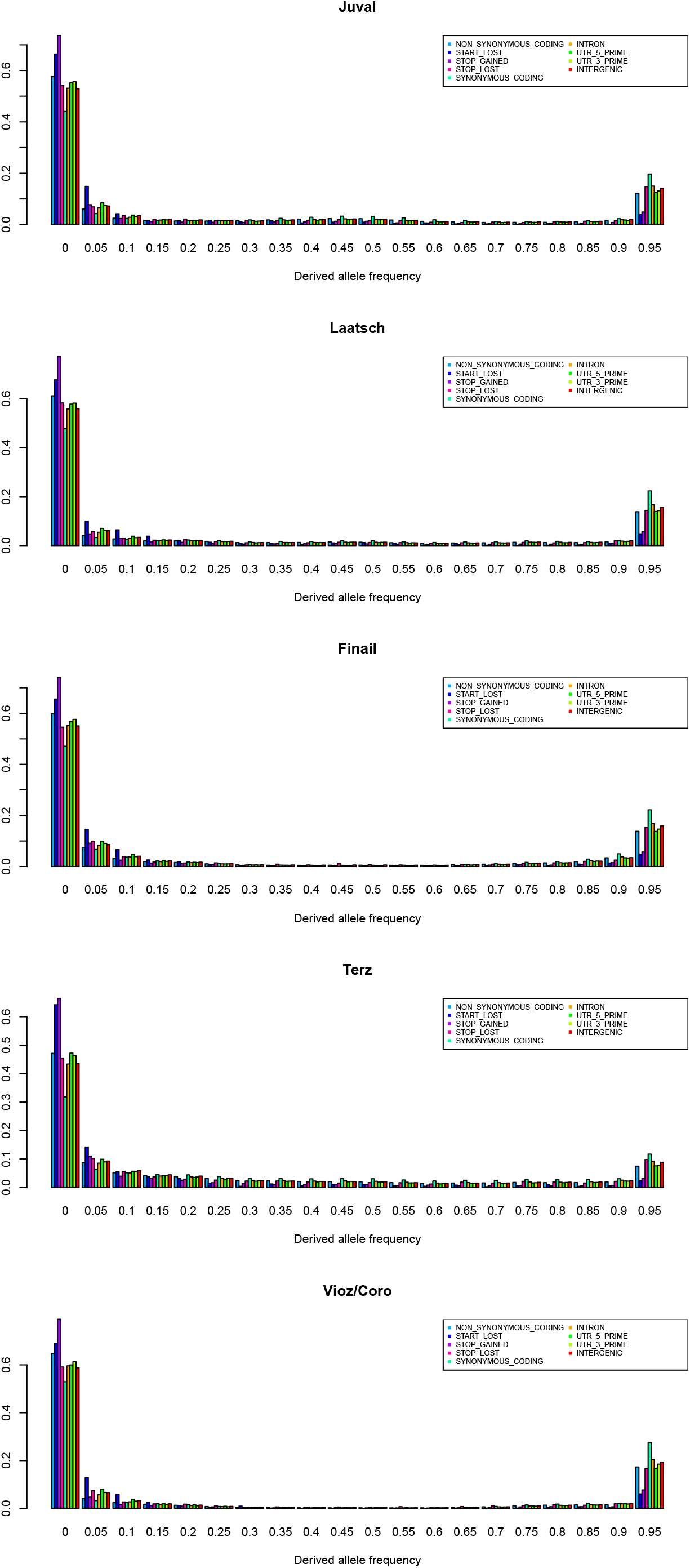
Derived allele frequency distributions across all populations for different SNP types

**Figure S5:**
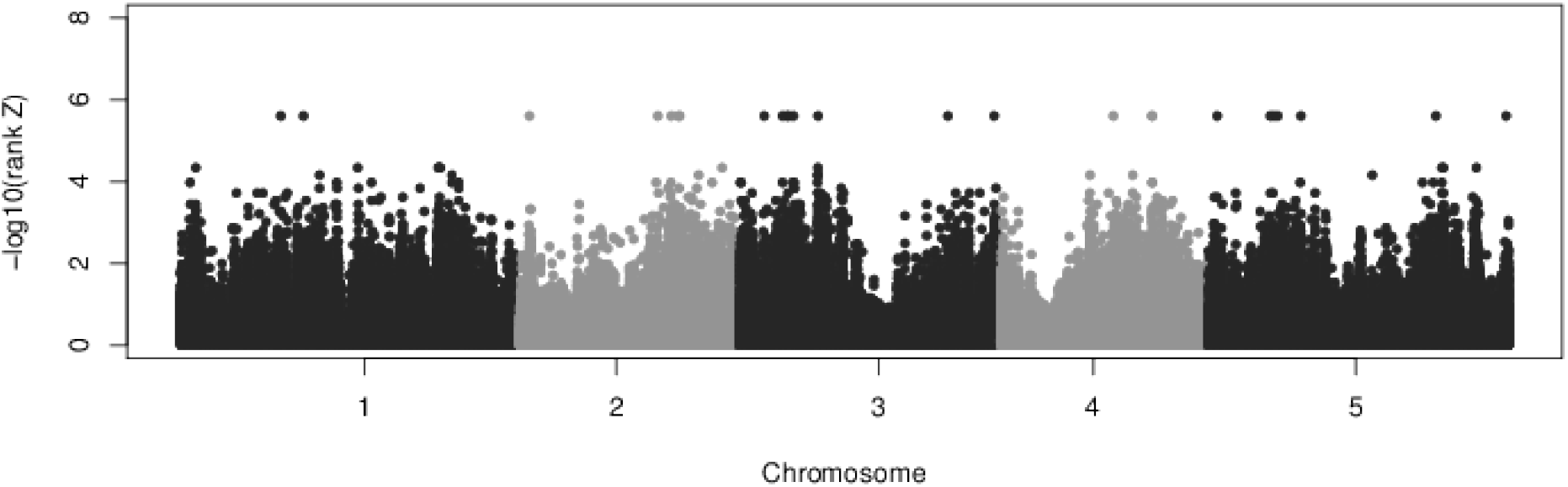
Manhattan plot of the Bayenv results.

**Figure S6:**
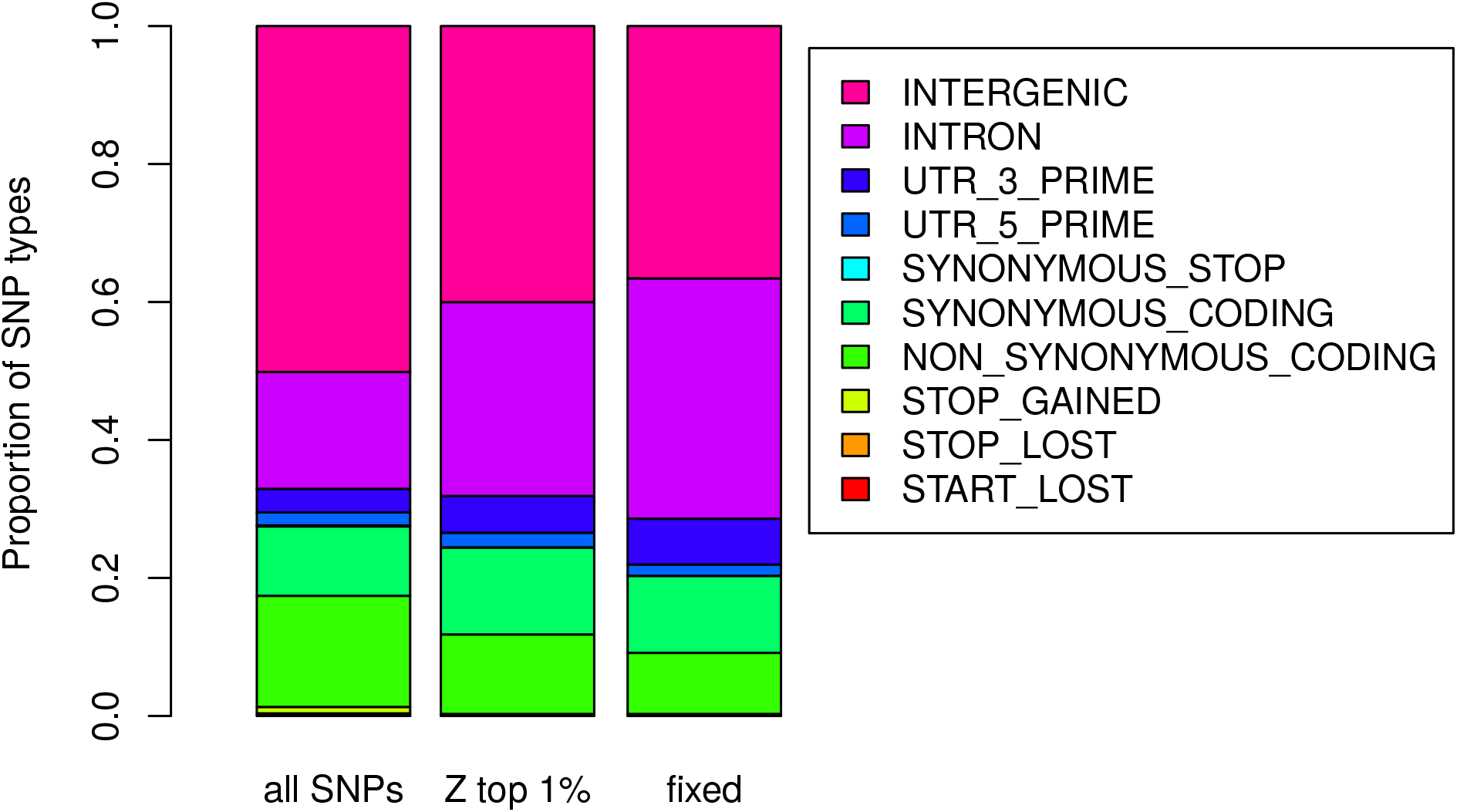
Relative proportion of the different SNP types for all SNPs and three different candidate SNP sets. (Bayenv2.0 top 1% and completely fixed between high and low pops)

**Figure S7:**
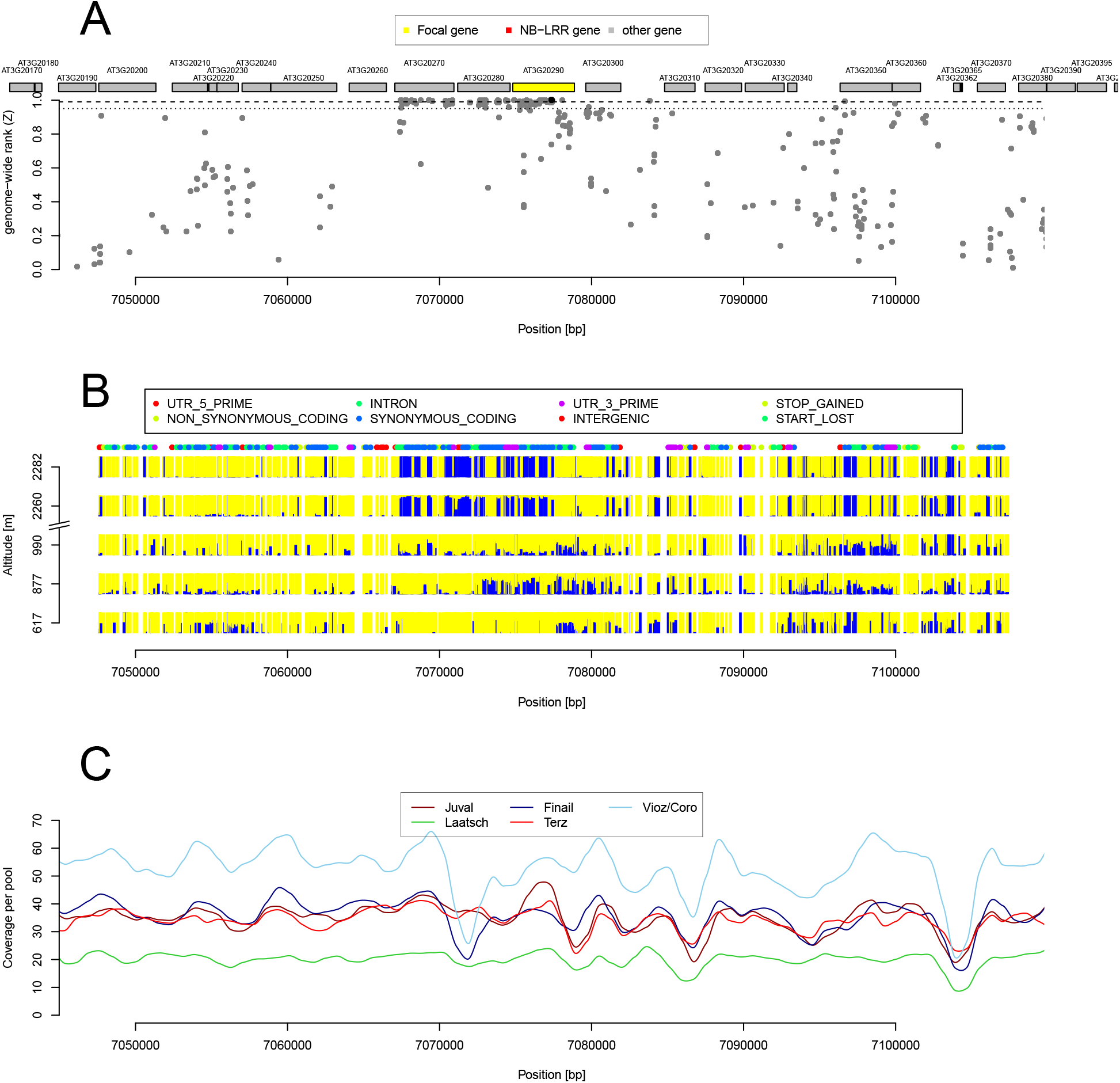
Example of the candidate gene AtEHD1 (AT3G20290) in which allele frequencies were highly differentiated between high and low altitude populations. (A) Plot of the genome-wide relative rank of the *Z* statistics. High values indicate a strong differentiation. (B) Plot of the relative allele frequencies of the two alleles segregating at a polymorphic site. The height of the bar is proportional to the allele frequency in the pooled sequence data. (C) Sequence coverage of the genomic region in the five different populations.

**Figure S8:**
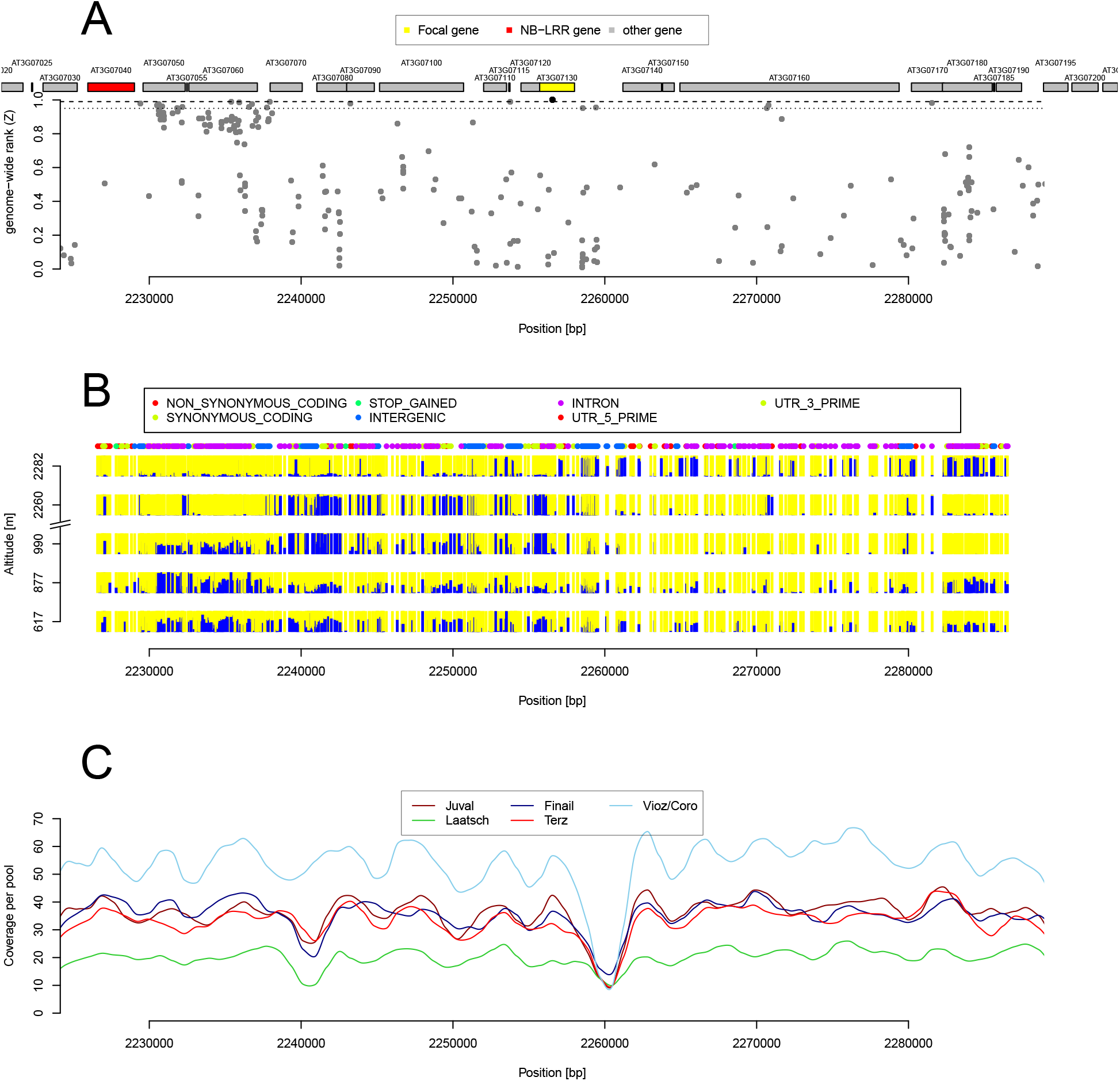
Example of a candidate gene (AT3G07130, encodes the purple acid phosphatase 15) in which allele frequencies differ strongly between the population at the highest altitude and the four lower altitude populations. (A) Plot of the genome-wide relative rank of the *Z* statistics. High values indicate a strong differentiation. (B) Plot of the relative allele frequencies of the two alleles segregating at a polymorphic site. The height of the par is proportional to the allele frequency. (C) Sequence coverage of the genomic region in the five different populations.

**Figure S9:**
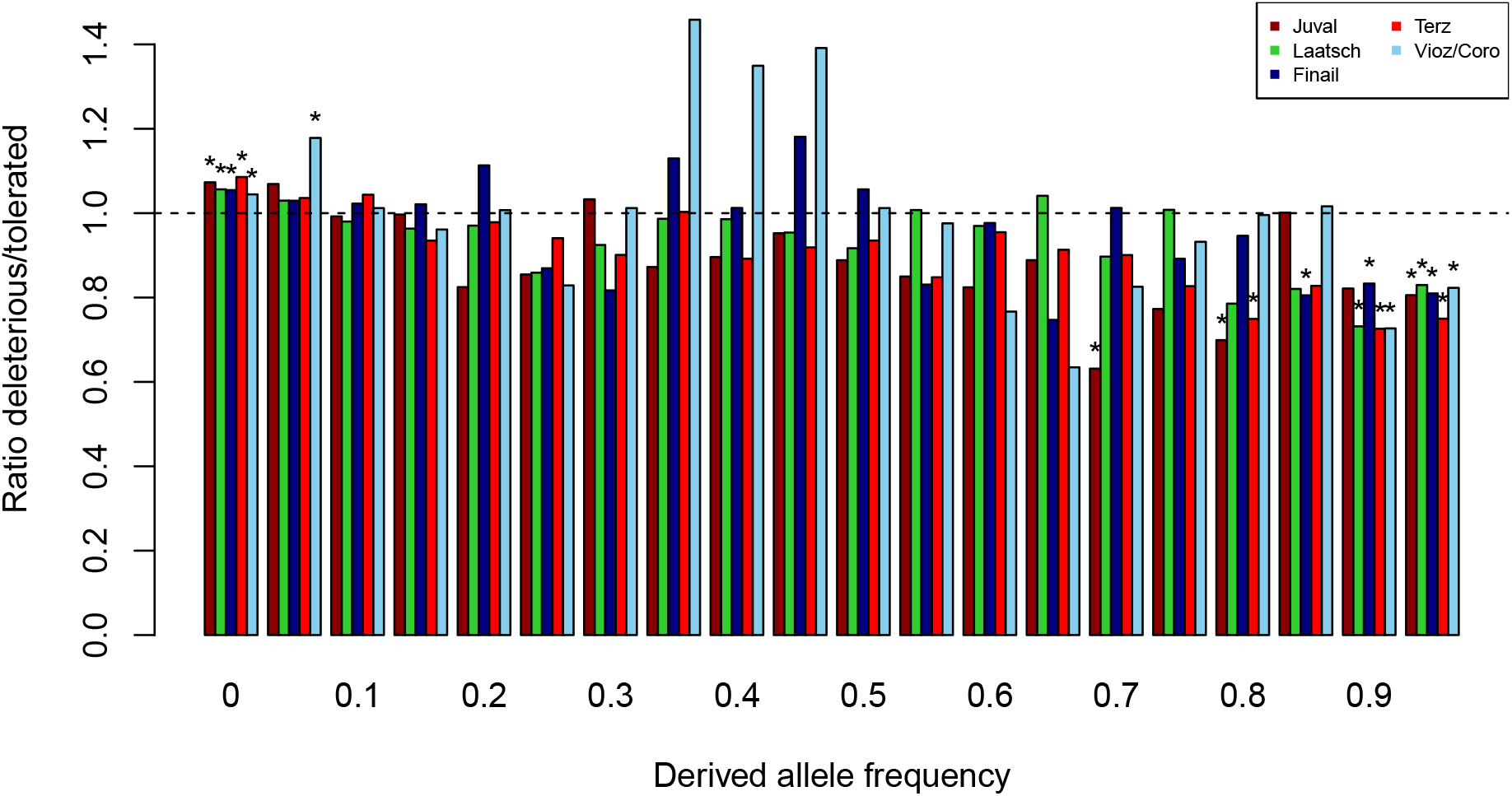
Ratio of relative frequencies of deleterious and tolerated amino acid polymorphisms per frequency bin. If both polymorphism types were neutral, a ratio of 1 (dashed line) would be expected for all frequency classes. Asterisk above bars represent ratios significantly different from 1 (nominal p-value threshold 0.01, tested with a χ^2^-test).

**Figure S10:**
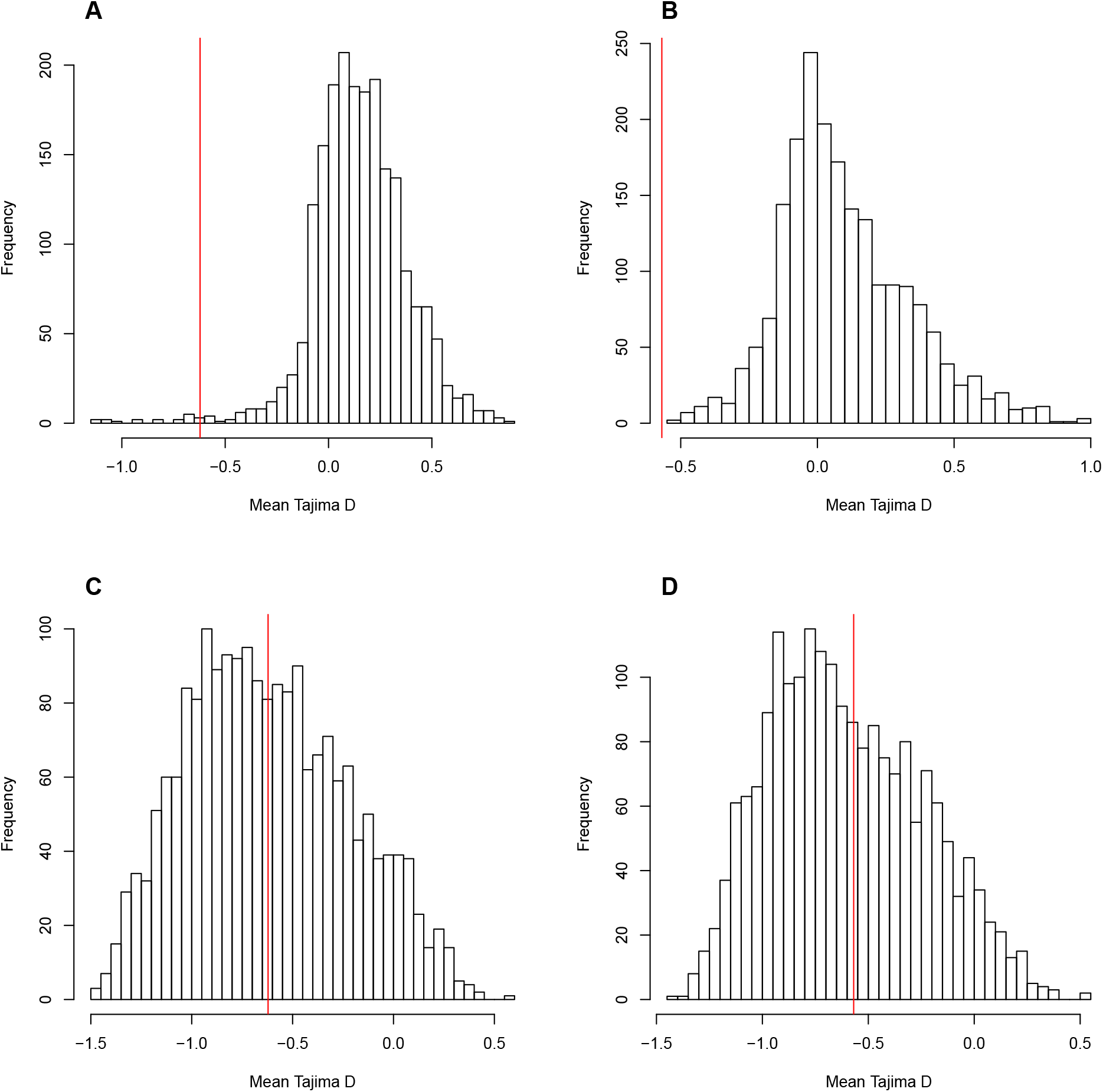
Posterior distribution of Tajima’s *D* values in the high altitude population for the best four ABC models. (A) Juval-Finail stepgrowth model. (B) Terz-Vioz/Coro exponential growth model. (C) Juval-Finail exponential growth with migration model. (D) Terz-Vioz/Coro exponential growth with migration model. Observed values are indicated by the red vertical lines.

### Supplementary Tables

**Table S1:**
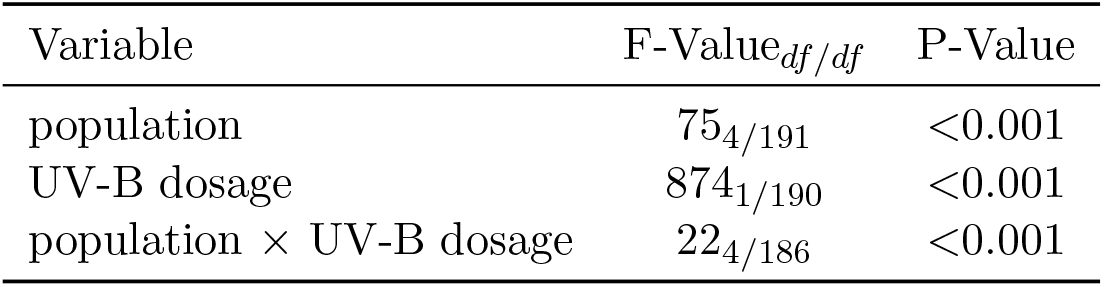
Analysis of deviance table of a binomial GLM for dead biomass in response to different dosages of UV-B.

**Table S2:** Average pairwise *F*_*ST*_ (lower triangle) and pairwise correlations of Bayenv2.0 correlation matrix among allele frequencies (upper triangle).

|  | Juval | Laatsch | Finail | Terz | Vioz/Coro |
| --- | --- | --- | --- | --- | --- |
| Juval | - | 0.79 | 0.81 | 0.75 | 0.67 |
| Laatsch | 0.12 | - | 0.77 | 0.99 | 0.71 |
| Finail | 0.12 | 0.12 | - | 0.71 | 0.81 |
| Terz | 0.12 | 0.12 | 0.14 | - | 0.62 |
| Vioz/Coro | 0.14 | 0.12 | 0.12 | 0.16 | - |

**Table S3:** Prior distributions of the Juval-Finail step-growth and Terz-Vioz/Coro exponential growth models.

| Parameter | J-F Stepgrowth |  | J-F Exponential mig. |  | T-V/C Exponential |  | T-V/C Exponential mig. |  |
| --- | --- | --- | --- | --- | --- | --- | --- | --- |
|  | Min | Max | Min | Max | Min | Max | Min | Max |
| $\theta^a$ | 2.8 | 11.2 | 2.8 | 11.2 | 2.8 | 11.2 | 2.8 | 11.2 |
| $\rho^b$ | 0.25 | 1.02 | 0.25 | 1.02 | 0.25 | 1.02 | 0.25 | 1.02 |
| $N_{high}$ | 0.001 | 1.0 | 0.001 | 1.0 | 0.001 | 1.0 | 0.001 | 1.0 |
| $N_{high,growth}^c$ | - | - | 0.0 | 159.7575 | 0.0 | 159.7575 | 0.0 | 159.7575 |
| $N_{high,growth,time}^d$ | - | - | 0.011875 | 0.4975 | 0.011875 | 0.4975 | 0.011875 | 0.4975 |
| $T_{split}$ | 0.0125 | 0.5 | 0.0125 | 0.5 | 0.0125 | 0.5 | 0.0125 | 0.5 |
| $M_{12}^e$ | - | - | 0.0 | 10.0 | - | - | 0.0 | 10.0 |
| $M_{21}$ | - | - | 0.0 | 10.0 | - | - | 0.0 | 10.0 |

<sup>a</sup>Theta is calculated as $4 * N_0 * \mu * L$ ; $N_0=[50,000:200,000]$ ; $\mu=7 * 10^{-9}$ ; $L=2000$ (length of simulated segments).
<sup>b</sup>Rho is calculated as $4 * N_0 * r * L$ ; $N_0=[50,000:200,000]$ ; $r=0.625 * 10^{-9}$ ; $L=2000$ (length of simulated segments).
<sup>c</sup>Growth rate of the high population.
<sup>d</sup>Time of the high population growth.
<sup>e</sup>Migration rate is calculated as $4 * N_0 * m$ .

**Table S4:** Posterior model probabilities of each demographic scenario calculated by the ABC *postpr* function using the *mnlogistic* method with a 0.005 tolerance to select the best growth model for the Juval-Finail and Terz-Vioz/Coro population pairs.

| Population pair | Without migration |  |  |  | With migration |  |  |  |
| --- | --- | --- | --- | --- | --- | --- | --- | --- |
|  | No growth | Step | Linear | Exponential | No growth | Step | Linear | Exponential |
| Juval-Finail | 0.000 | 0.00 | 0.000 | 0.000 | 0.000 | 0.001 | 0.000 | 0.999 |
| Terz-Vioz/Coro | 0.000 | 0.000 | 0.000 | 0.000 | 0.000 | 0.000 | 0.000 | 1.000 |

**Table S5:** Soil parameters of surface soil (5-10 cm) in the five population sites.

| Population | pH | Organic matter<br>% | Phosphate<br>mg/100g | Potassium<br>mg/100g | Magnesium<br>mg/100g |
| --- | --- | --- | --- | --- | --- |
| Juval | 6.1 | 6.17 | 4.2 | 21 | 21 |
| Terz | 6.2 | 4.9 | 5.5 | 29 | 26 |
| Laatsch | 6.3 | 3.47 | <1.0 | 8 | 8.3 |
| Finail | 4.6 | 17.2 | 16 | 58 | 36 |
| Vioz/Coro | 5.2 | 15.8 | 22 | 24 | 23 |

**Table S6:** All 25 SNPs with *Z* = 0.5 in the Bayenv2 analysis.

| Chromosome | bp | AGI ID | SNP type | Description |
| --- | --- | --- | --- | --- |
| 1 | 9,021,128 | AT1G26090 | Intron | P-loop containing nucleoside triphosphate hydrolases superfamily protein |
| 1 | 11,056,544 | AT1G31010 | Synonymous coding | Organellar single-stranded DNA binding protein 4 (OSB4) |
| 2 | 902,758 | AT2G03060 | Intron | *AGAMOUS-LIKE 30*; Pollen development, pollen exine formation |
| 2 | 12,412,973 |  | Intergenic |  |
| 2 | 13,606,764 | AT2G31970 | Non-synonymous coding | Encodes the Arabidopsis RAD50 homologue; DNA repair, RNA processing |
| 2 | 14,276,102 | AT2G33760 | Synonymous coding | Pentatricopeptide repeat (PPR) superfamily protein |
| 2 | 14,376,598 | AT2G34030 | Intron | Calcium-binding EF-hand family protein |
| 3 | 2,256,537 | AT3G07130 | Non-synonymous coding | *PURPLE ACID PHOSPHATASE 15*; Pollen germination, seed germination |
| 3 | 3,897,644 | AT3G12220 | Intron | *SERINE CARBOXYPEPTIDASE-LIKE 16*; Proteolysis |
| 3 | 4,298,285 | AT3G13290 | Non-synonymous coding | Varicose-related (*VCR*) |
| 3 | 4,427,242 | AT3G13560 | Non-synonymous coding | O-Glycosyl hydrolases family 17 protein; Carbohydrate metabolic process |
| 3 | 4,878,614 |  | Intergenic |  |
| 3 | 7,077,355 | AT3G20290 | Intron | Arabidopsis Eps15 homology domain proteins involved in endocytosis (*AtEHD1*) |
| 3 | 18,716,502 | AT3G50430 | Non-synonymous coding | Unknown protein |
| 3 | 22,868,343 | AT3G61780 | Non-synonymous coding | Embryo defective 1703 (emb1703); Embryo development ending in seed dormancy |
| 4 | 10,106,954 |  | Intergenic |  |
| 4 | 13,574,591 | AT4G27040 | 5' UTR | VPS22; Vesicle-mediated transport |
| 4 | 13,577,662 | AT4G27050 | Intron | F-box/RNI-like superfamily protein |
| 5 | 824,047 | AT5G03360 | Intron | DC1 domain-containing protein; Oxidation-reduction process |
| 5 | 5,587,807 | AT5G17010 | Intron | Major facilitator superfamily protein; Transmembrane transport |
| 5 | 5,903,750 | AT5G17860 | Non-synonymous coding | Calcium exchanger 7 (*CAX7*) |
| 5 | 6,252,514 |  | Intergenic |  |
| 5 | 8,335,327 |  | Intergenic |  |
| 5 | 20,429,419 |  | Intergenic |  |
| 5 | 26,727,336 | AT5G66930 | Intron | Unknown protein |

**Table S7:**
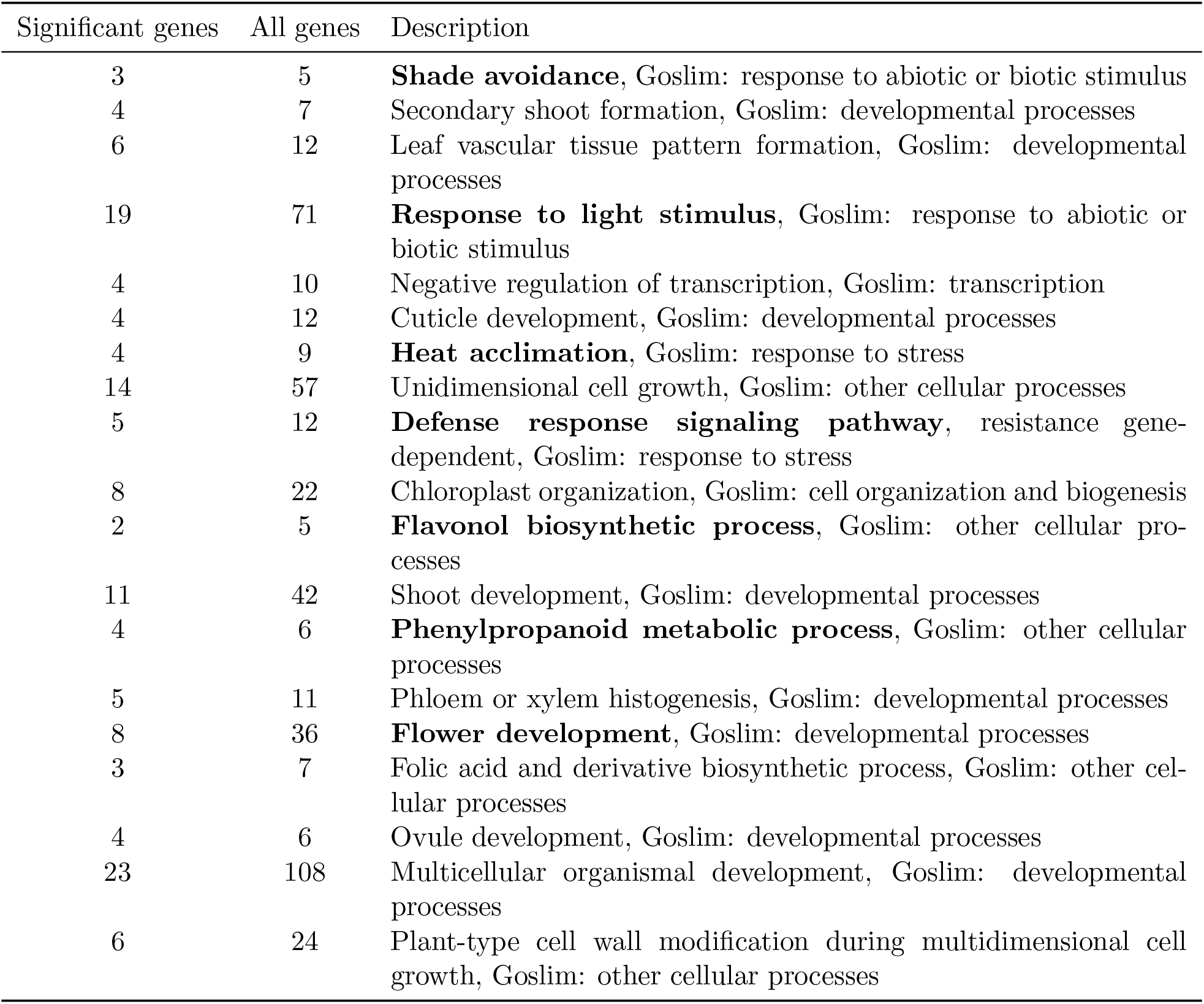
Enriched GO Biological processes among the top Bayenv2.0 results with nominal p-values ≤ 0.05.

**Table S8:** Enriched PO, KEGG, AraCyc categories among the top Bayenv2.0 results with nominal p-values ≤ 0.05.

| Significant genes | All genes | Description |
| --- | --- | --- |
| 9 | 15 | PO:0020144 expressed in apical meristem |
| 14 | 38 | PO:0000229 **expressed in flower meristem** |
| 3 | 4 | **Superoxide radicals degradation** |
| 6 | 12 | **Glucosinolate biosynthesis** |
| 8 | 20 | CDP-diacylglycerol biosynthesis I |
| 8 | 20 | CDP-diacylglycerol biosynthesis II |
| 5 | 10 | glycolipid desaturation |
| 5 | 11 | PO:0020149 expressed in quiescent center |
| 4 | 9 | Glucosinolate biosynthesis from phenylalanine |
| 3 | 6 | Glucosinolate biosynthesis from tetrahomomethionine |
| 3 | 6 | Glucosinolate biosynthesis from trihomomethionine |
| 5 | 11 | Tetrahydrofolate biosynthesis II |
| 10 | 34 | Protein export |
| 3 | 5 | Mevalonate pathway I |
| 3 | 7 | Glucosinolate biosynthesis from homomethionine |
| 3 | 7 | Glucosinolate biosynthesis from hexahomomethionine |
| 3 | 7 | Glucosinolate biosynthesis from pentahomomethionine |
| 2 | 3 | Brassinosteroid biosynthesis II |
| 7 | 22 | PO:0006085 expressed in root meristem |
| 2 | 2 | PO:0007130 expressed during B reproductive growth |
| 3 | 7 | Glucosinolate biosynthesis from dihomomethionine |
| 8 | 23 | α-Linolenic acid metabolism |
| 3 | 4 | **Flavone and flavonol biosynthesis** |
| 4 | 8 | Isoleucine degradation I |
| 3 | 3 | **Anthocyanin biosynthesis** (pelargonidin 3-O-glucoside, cyanidin 3-O-glucoside) |
| 3 | 8 | PO:0000372 expressed in metaxylem |
| 7 | 19 | FormylTHF biosynthesis II |

**Table S9:** GO terms among top *Z* signals with nominal p-value≤ 0.05 obtained from an analysis with Gowinda.

| Cat | FDR | Genes found | All genes | Description |
| --- | --- | --- | --- | --- |
| CC | 0.2918 | 29 | 125 | GO:0009579 thylakoid, Goslim:other intracellular components |
| MF | 0.30478875 | 3 | 4 | GO:0004784 superoxide dismutase activity, GOslim other enzyme activity |
| MF | 0.30478875 | 7 | 14 | GO:0008794 arsenate reductase (glutaredoxin) activity, GOslim other enzyme activity |
| BP | 0.30478875 | 2 | 2 | GO:0016579 protein deubiquitination, Goslim:protein metabolism |
| BP | 0.30478875 | 3 | 5 | GO:0009641 shade avoidance, Goslim:response to abiotic or biotic stimulus |
| CC | 0.30478875 | 9 | 22 | GO:0009295 nucleoid, Goslim:other intracellular components |
| CC | 0.30478875 | 5 | 10 | GO:0005819 spindle, Goslim:other intracellular components |
| MF | 0.30478875 | 208 | 1185 | GO:0003700 transcription factor activity, GOslim transcription factor activity |
| BP | 0.3569411111 | 4 | 7 | GO:0010223 secondary shoot formation, Goslim:developmental processes |
| CC | 0.3764346154 | 6 | 15 | GO:0019005 SCF ubiquitin ligase complex, Goslim:other intracellular components |
| CC | 0.3764346154 | 50 | 221 | GO:0005783 endoplasmic reticulum, Goslim:ER |
| BP | 0.3764346154 | 6 | 12 | GO:0010305 leaf vascular tissue pattern formation, Goslim:developmental processes |
| MF | 0.3764346154 | 17 | 58 | GO:0046872 metal ion binding, GOslim other binding |
| CC | 0.3783357143 | 8 | 21 | GO:0009524 phragmoplast, Goslim:other cytoplasmic components |
| CC | 0.44608 | 3 | 5 | GO:0009574 preprophase band, Goslim:other cytoplasmic components |
| BP | 0.493291875 | 19 | 71 | GO:0009416 response to light stimulus, Goslim:response to abiotic or biotic stimulus |
| MF | 0.5177496 | 13 | 56 | GO:0016563 transcription activator activity, GOslim other molecular functions |
| BP | 0.5177496 | 4 | 10 | GO:0016481 negative regulation of transcription, Goslim:transcription |
| BP | 0.5177496 | 4 | 12 | GO:0042335 cuticle development, Goslim:developmental processes |
| BP | 0.5177496 | 4 | 9 | GO:0010286 heat acclimation, Goslim:response to stress |
| BP | 0.5177496 | 14 | 57 | GO:0009826 unidimensional cell growth, Goslim:other cellular processes |
| BP | 0.5177496 | 5 | 12 | GO:0009870 defense response signaling pathway, resistance gene-dependent, Goslim:response to stress |
| BP | 0.5177496 | 8 | 22 | GO:0009658 chloroplast organization, Goslim:cell organization and biogenesis |
| MF | 0.5177496 | 6 | 15 | GO:0008017 microtubule binding, GOslim protein binding |
| BP | 0.5177496 | 2 | 5 | GO:0051555 flavonol biosynthetic process, Goslim:other cellular processes |
| MF | 0.5183137037 | 13 | 48 | GO:0008415 acyltransferase activity, GOslim transferase activity |
| BP | 0.5183137037 | 11 | 42 | GO:0048367 shoot development, Goslim:developmental processes |
| BP | 0.5257717857 | 4 | 6 | GO:0009698 phenylpropanoid metabolic process, Goslim:other cellular processes |
| BP | 0.5337251724 | 5 | 11 | GO:0010087 phloem or xylem histogenesis, Goslim:developmental processes |
| BP | 0.5614648649 | 8 | 36 | GO:0009908 flower development, Goslim:developmental processes |
| MF | 0.5614648649 | 5 | 10 | GO:0016207 4-coumarate-CoA ligase activity, GOslim other enzyme activity |
| BP | 0.5614648649 | 3 | 7 | GO:0009396 folic acid and derivative biosynthetic process, Goslim:other cellular processes |
| MF | 0.5614648649 | 5 | 12 | GO:0016291 acyl-CoA thioesterase activity, GOslim hydrolase activity |
| BP | 0.5614648649 | 4 | 6 | GO:0048481 ovule development, Goslim:developmental processes |
| BP | 0.5614648649 | 23 | 108 | GO:0007275 multicellular organismal development, Goslim:developmental processes |
| CC | 0.5614648649 | 148 | 835 | GO:0005634 nucleus, Goslim:nucleus |
| BP | 0.5614648649 | 6 | 24 | GO:0009831 plant-type cell wall modification during multidimensional cell growth, Goslim:other cellular processes |
| CC | 0.5720152632 | 2 | 4 | GO:0016272 prefoldin complex, Goslim:cytosol |
| MF | 0.5730625641 | 46 | 205 | GO:0005215 transporter activity, GOslim transporter activity |

**Table S10:** KEGG pathways, PO terms and Aracyc networks among top *Z* signals with nominal p-value≤ 0.05

| Cat | FDR | Genes found | All genes | Description |
| --- | --- | --- | --- | --- |
| PO | 0.0746 | 9 | 15 | PO:0020144 expressed in apical meristem |
| PO | 0.133415 | 14 | 38 | PO:0000229 expressed in flower meristem |
| ARACYC | 0.1537485714 | 3 | 4 | superoxide radicals degradation |
| KEGG | 0.1537485714 | 6 | 12 | Glucosinolate biosynthesis |
| ARACYC | 0.1537485714 | 8 | 20 | CDP-diacylglycerol biosynthesis I |
| ARACYC | 0.1537485714 | 8 | 20 | CDP-diacylglycerol biosynthesis II |
| ARACYC | 0.1537485714 | 5 | 10 | glycolipid desaturation |
| PO | 0.1630625 | 5 | 11 | PO:0020149 expressed in quiescent center |
| ARACYC | 0.235128 | 4 | 9 | glucosinolate biosynthesis from phenylalanine |
| ARACYC | 0.2970092857 | 3 | 6 | glucosinolate biosynthesis from tetrahomomethionine |
| ARACYC | 0.2970092857 | 3 | 6 | glucosinolate biosynthesis from trihomomethionine |
| ARACYC | 0.2970092857 | 5 | 11 | tetrahydrofolate biosynthesis II |
| KEGG | 0.2970092857 | 10 | 34 | Protein export |
| ARACYC | 0.324188 | 3 | 5 | mevalonate pathway I |
| ARACYC | 0.3547026316 | 3 | 7 | glucosinolate biosynthesis from homomethionine |
| ARACYC | 0.3547026316 | 3 | 7 | glucosinolate biosynthesis from hexahomomethionine |
| ARACYC | 0.3547026316 | 3 | 7 | glucosinolate biosynthesis from pentahomomethionine |
| ARACYC | 0.3547026316 | 2 | 3 | brassinosteroid biosynthesis II |
| PO | 0.3764761905 | 7 | 22 | PO:0006085 expressed in root meristem |
| PO | 0.3764761905 | 2 | 2 | PO:0007130 expressed during B reproductive growth |
| ARACYC | 0.4155586957 | 3 | 7 | glucosinolate biosynthesis from dihomomethionine |
| KEGG | 0.4155586957 | 8 | 23 | alpha-Linolenic acid metabolism |
| KEGG | 0.42463625 | 3 | 4 | Flavone and flavonol biosynthesis |
| ARACYC | 0.4529372 | 4 | 8 | isoleucine degradation I |
| ARACYC | 0.4808680769 | 3 | 3 | anthocyanin biosynthesis (pelargonidin 3-O-glucoside, cyanidin 3-O-glucoside) |
| PO | 0.4842432609 | 3 | 8 | PO:0000372 expressed in metaxylem |
| ARACYC | 0.4842432609 | 7 | 19 | formylTHF biosynthesis II |

**Table S11:**
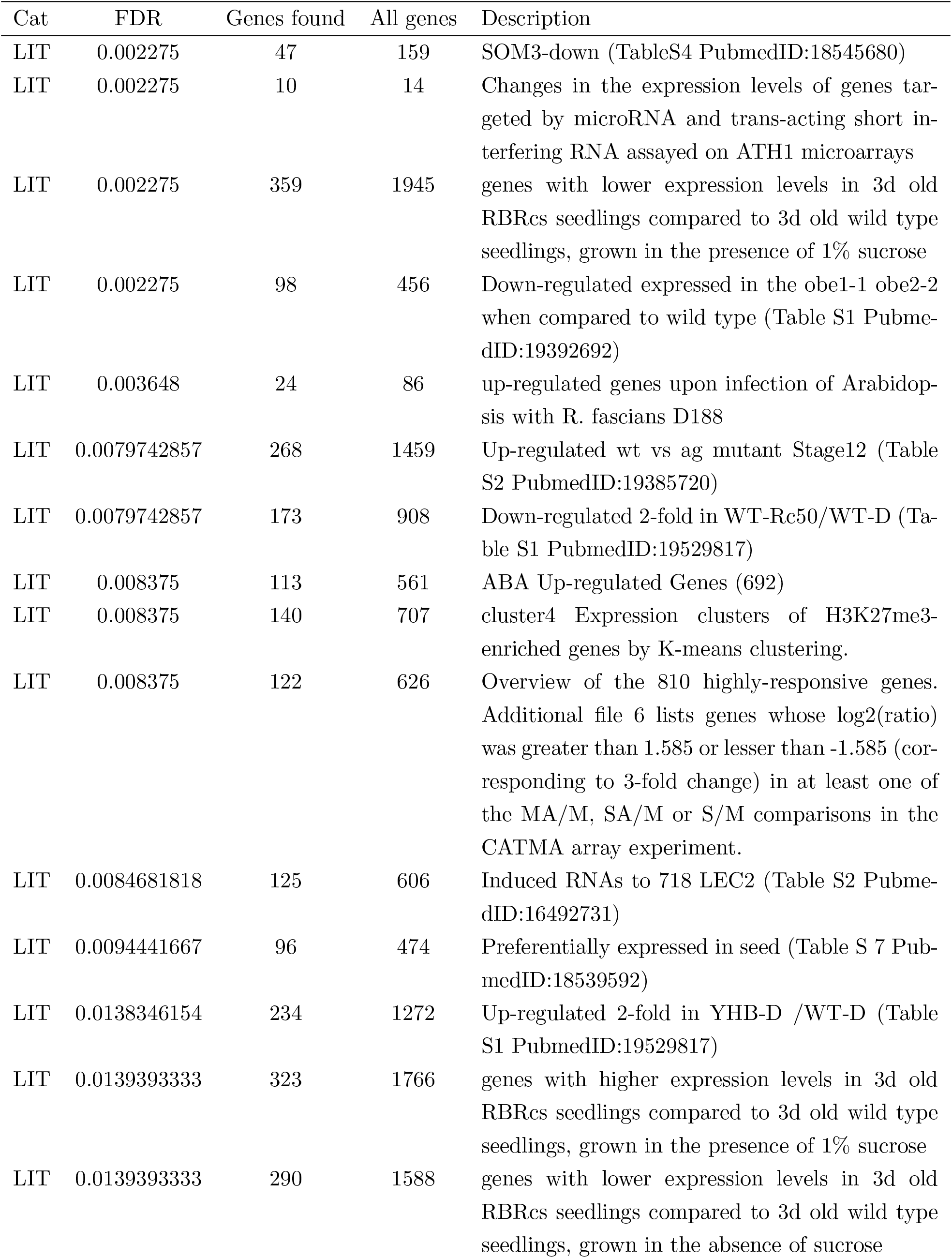
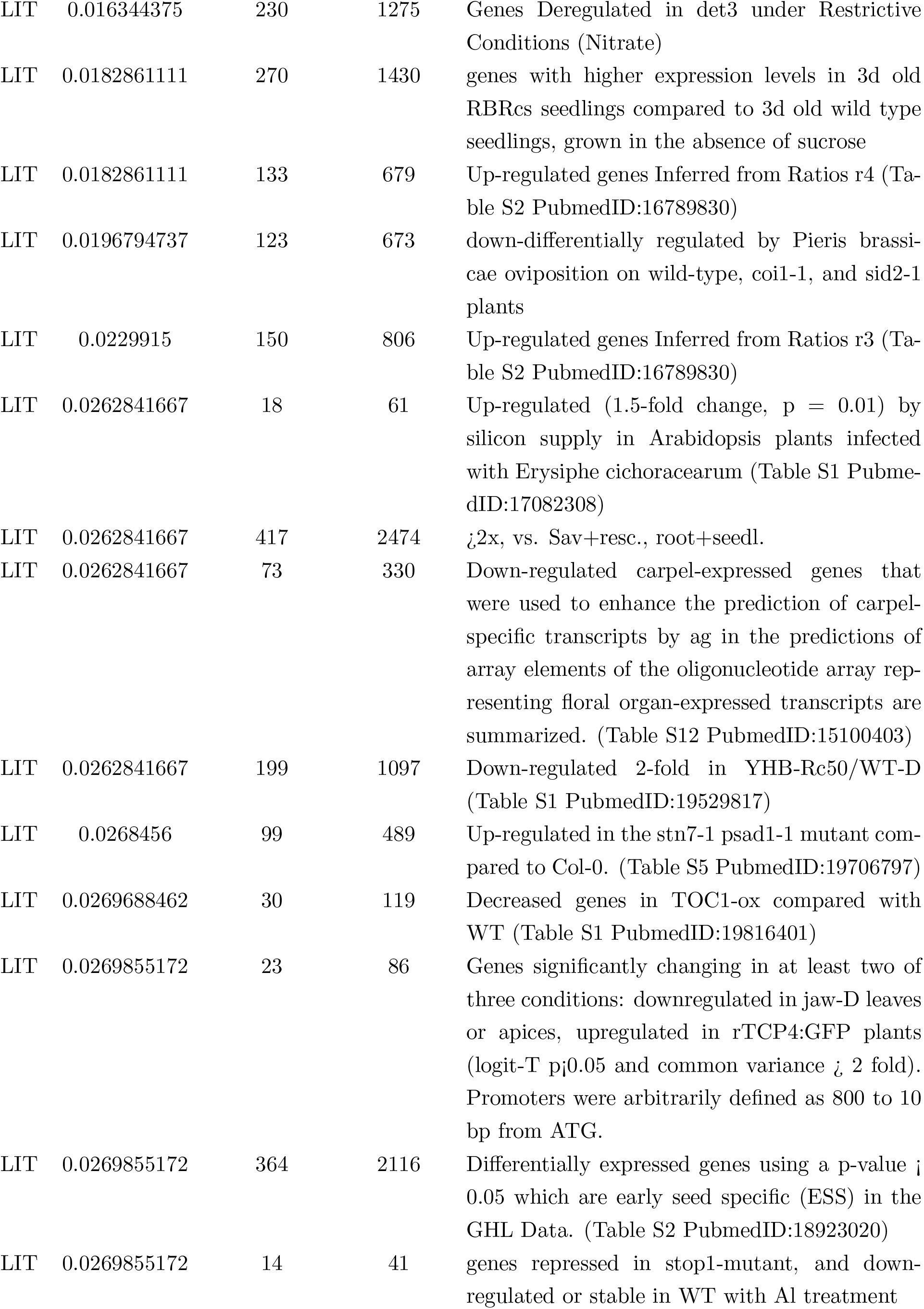
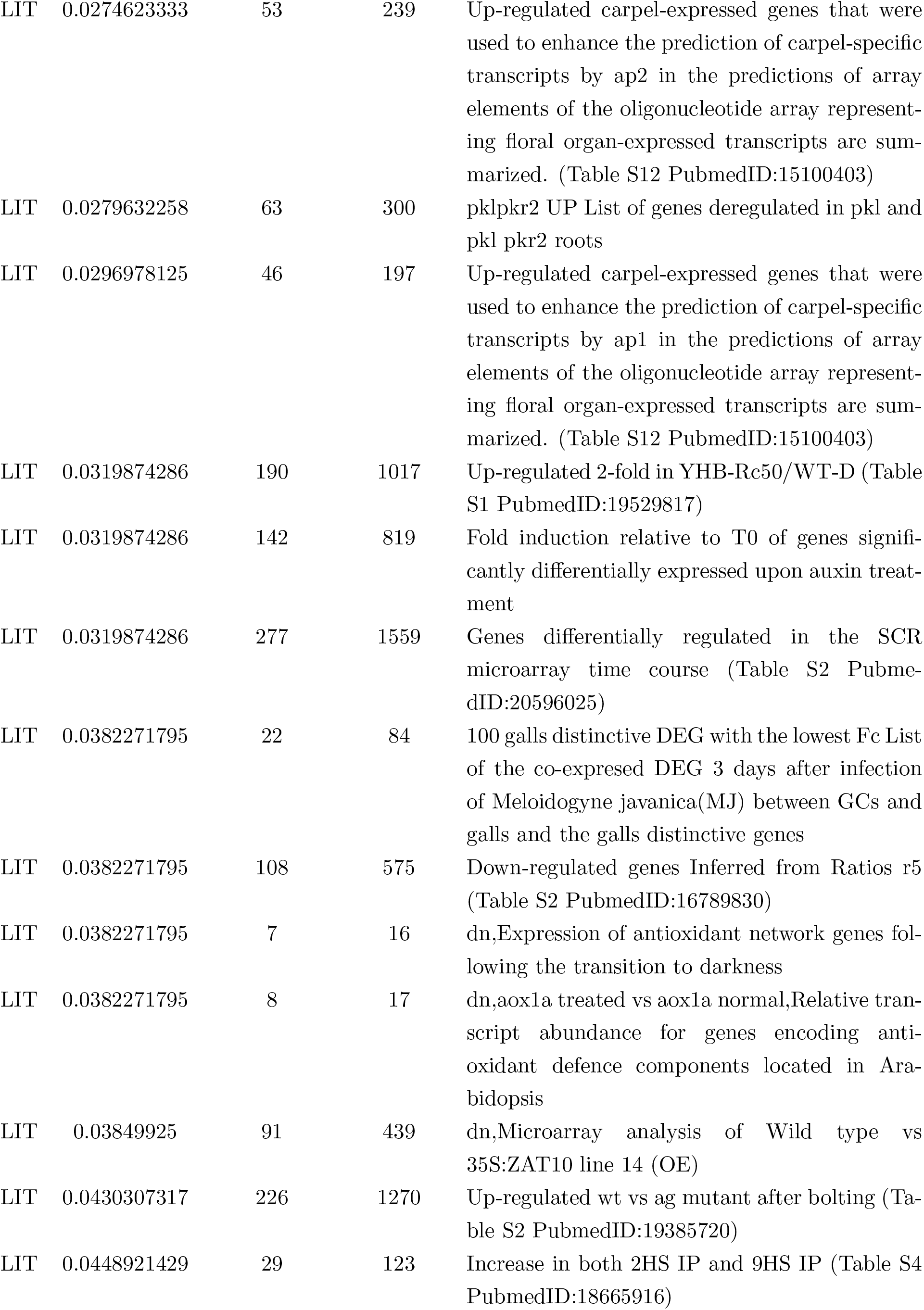
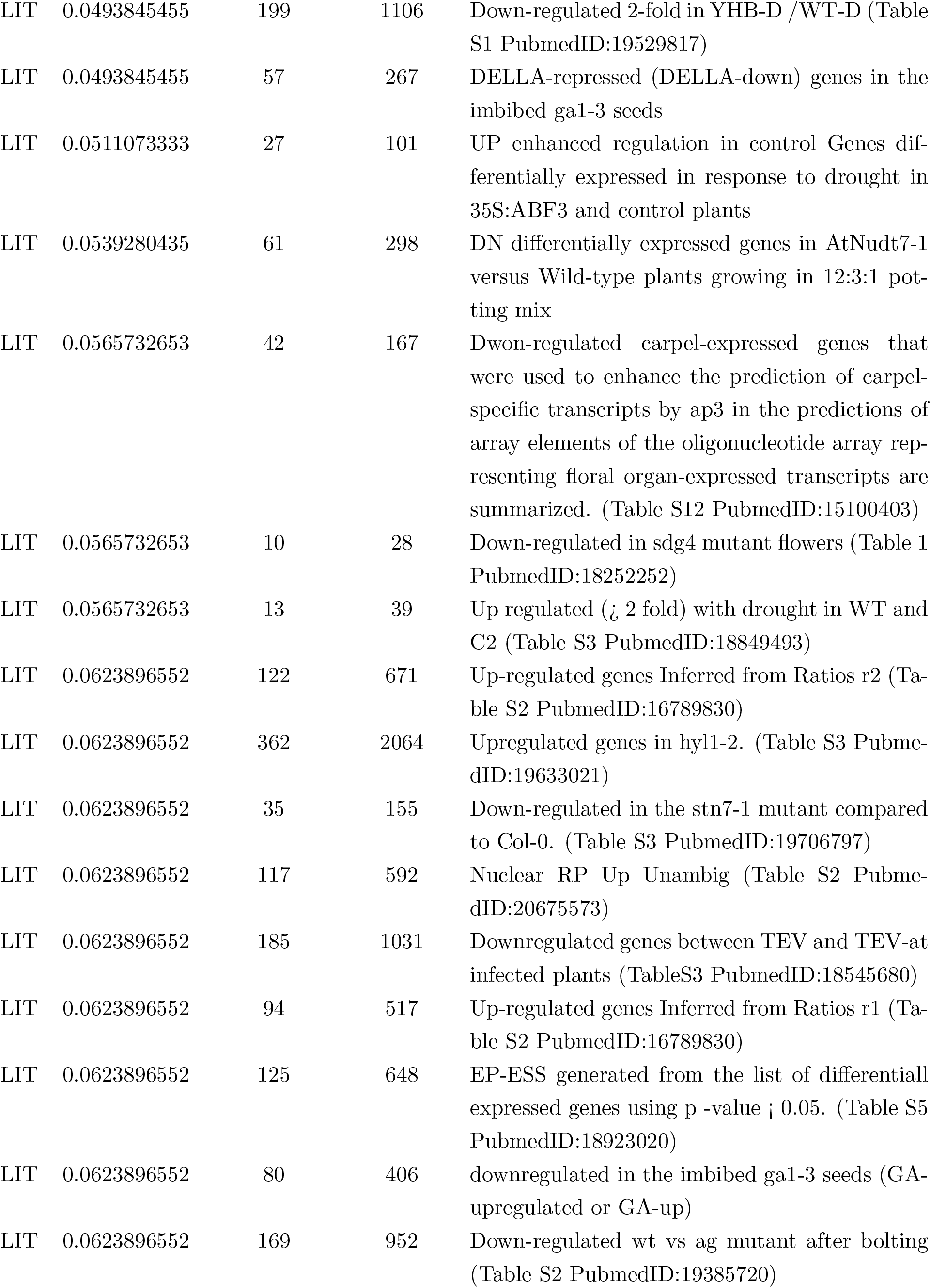
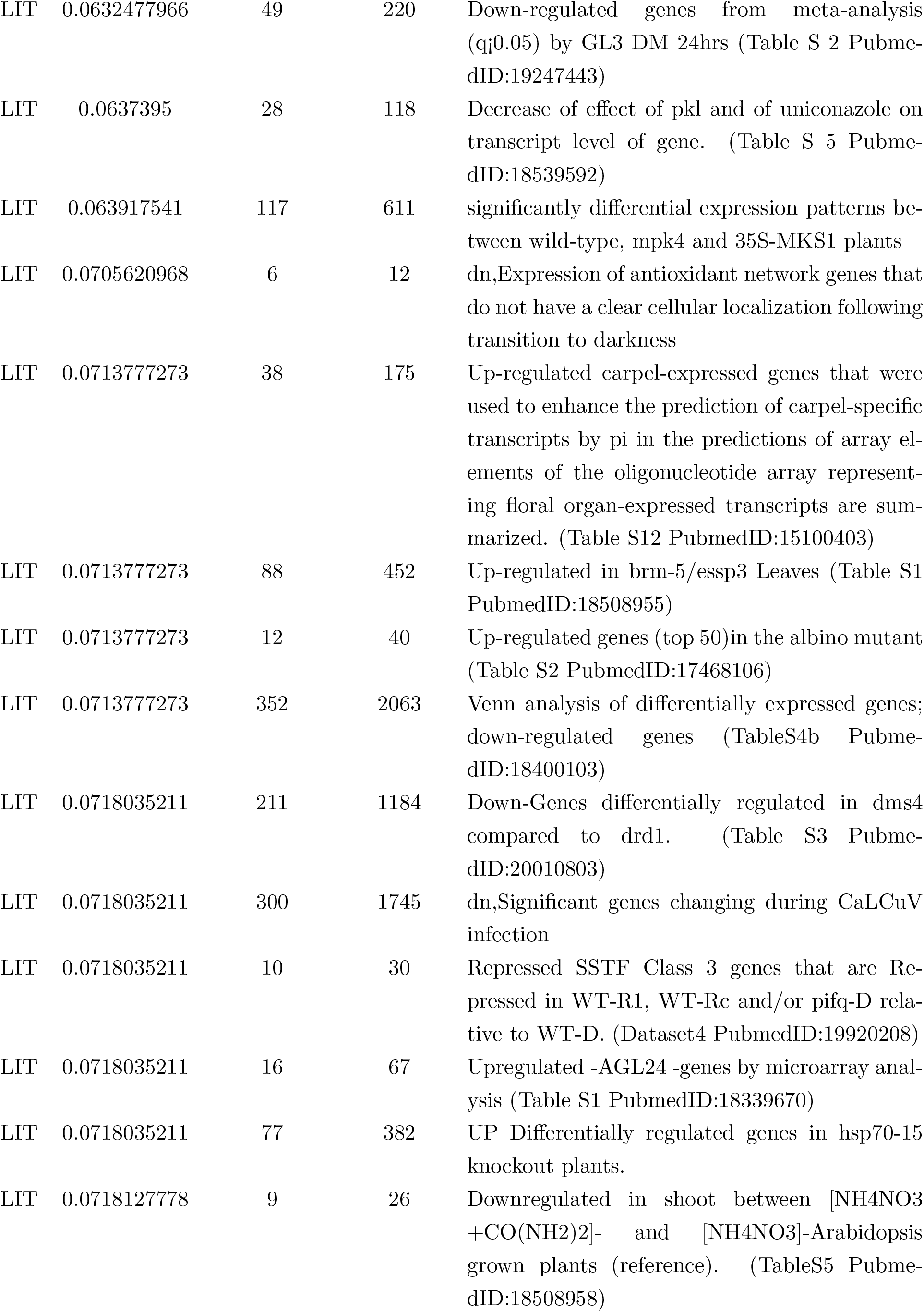
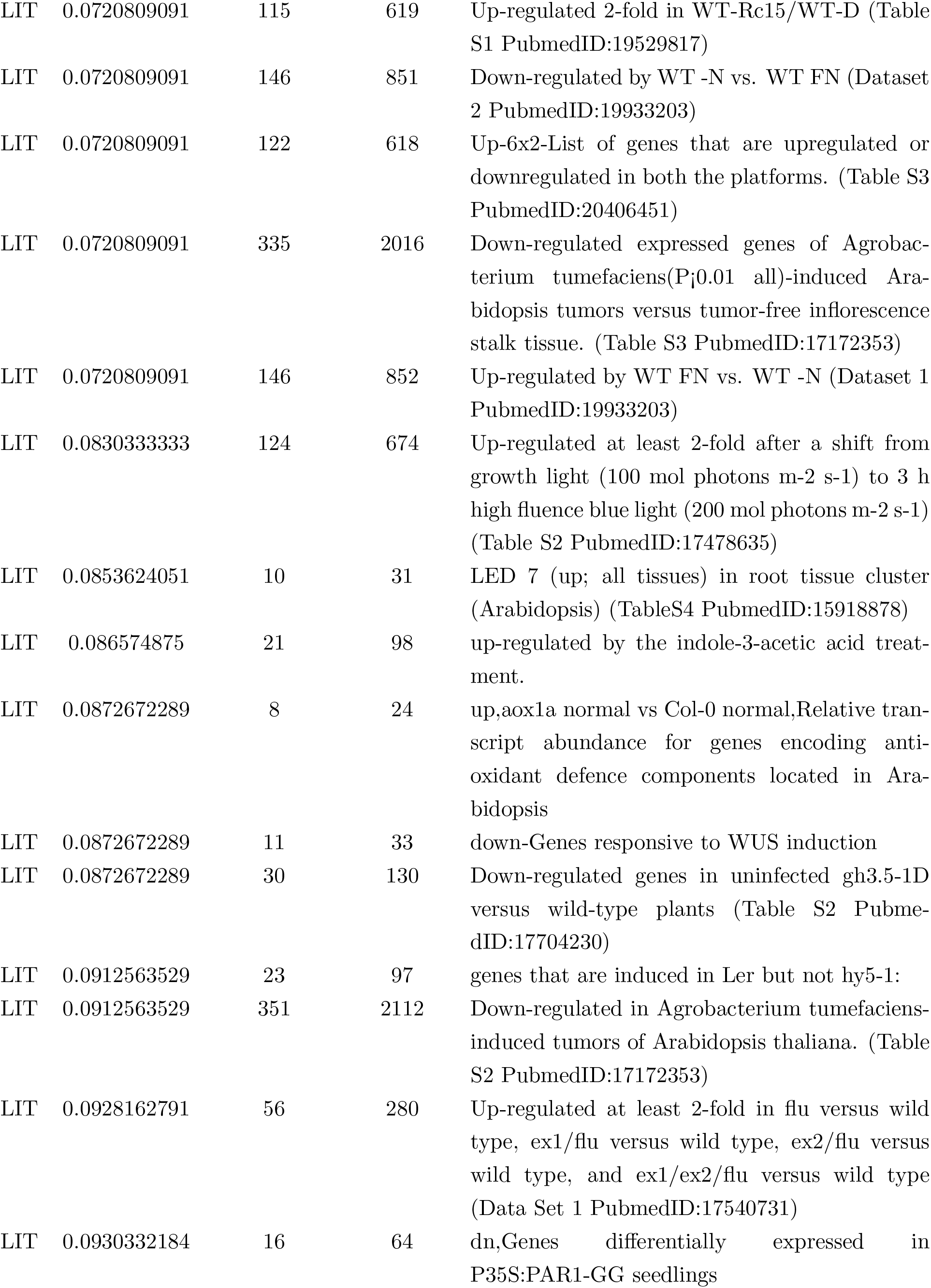
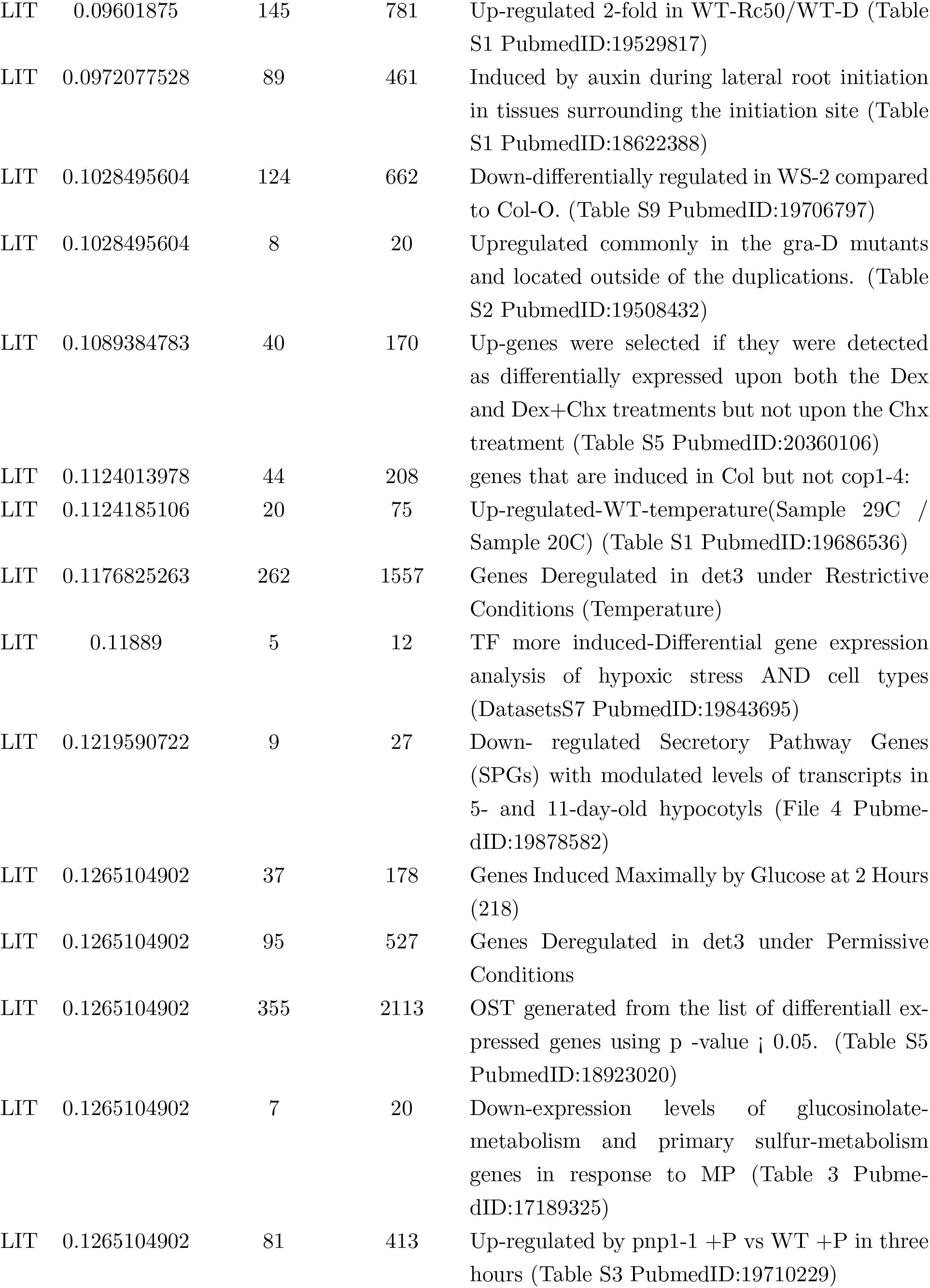
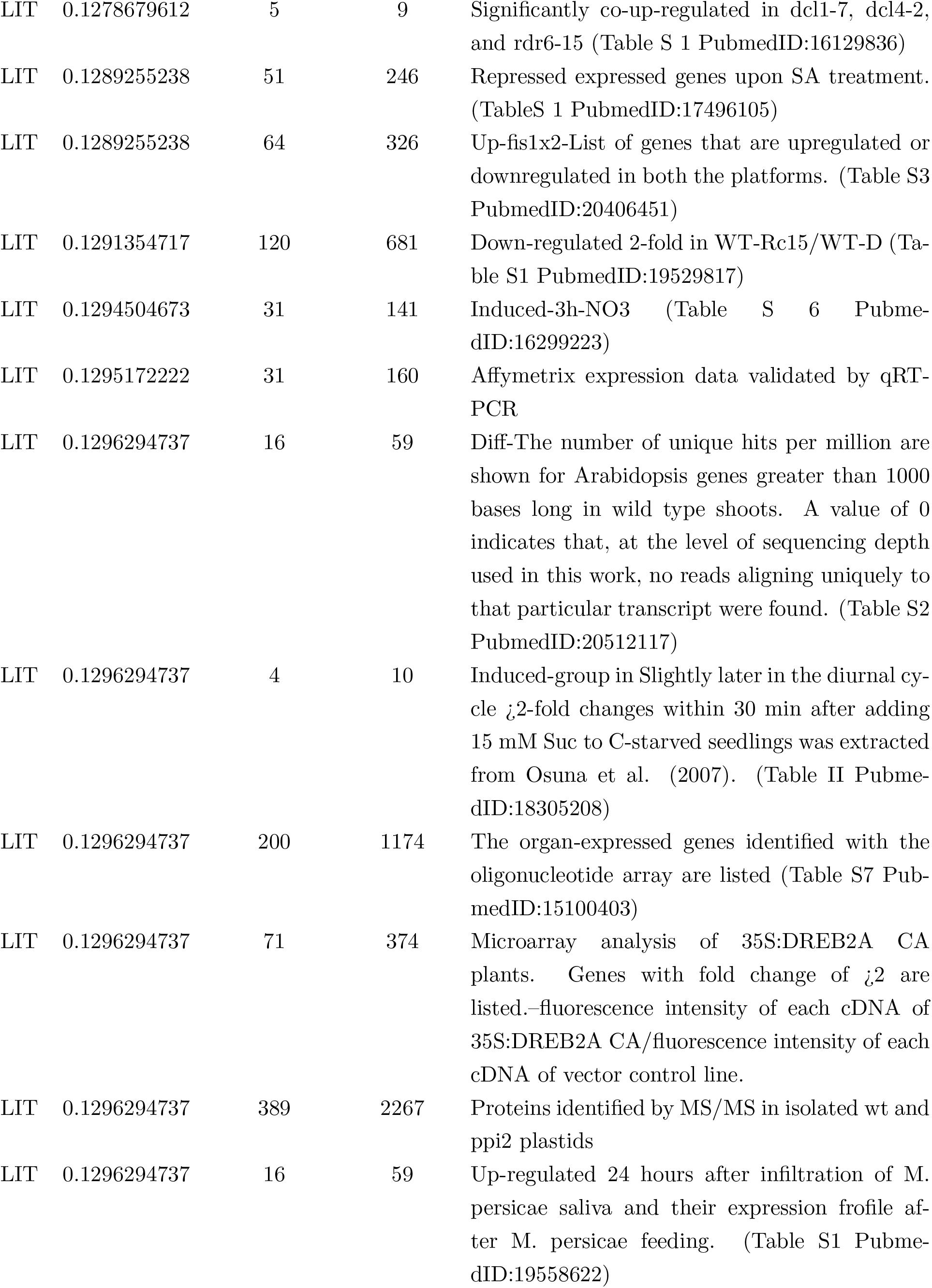
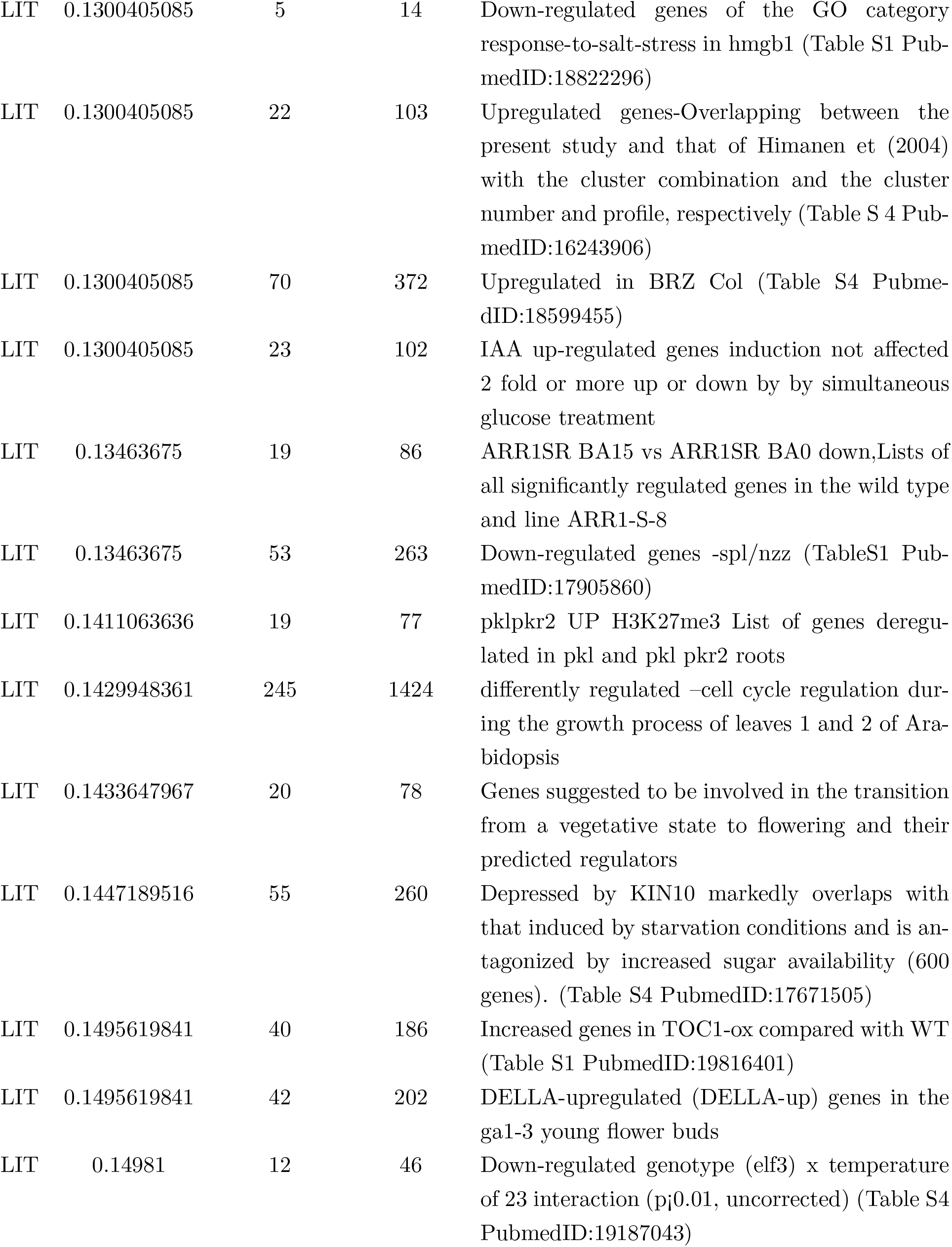
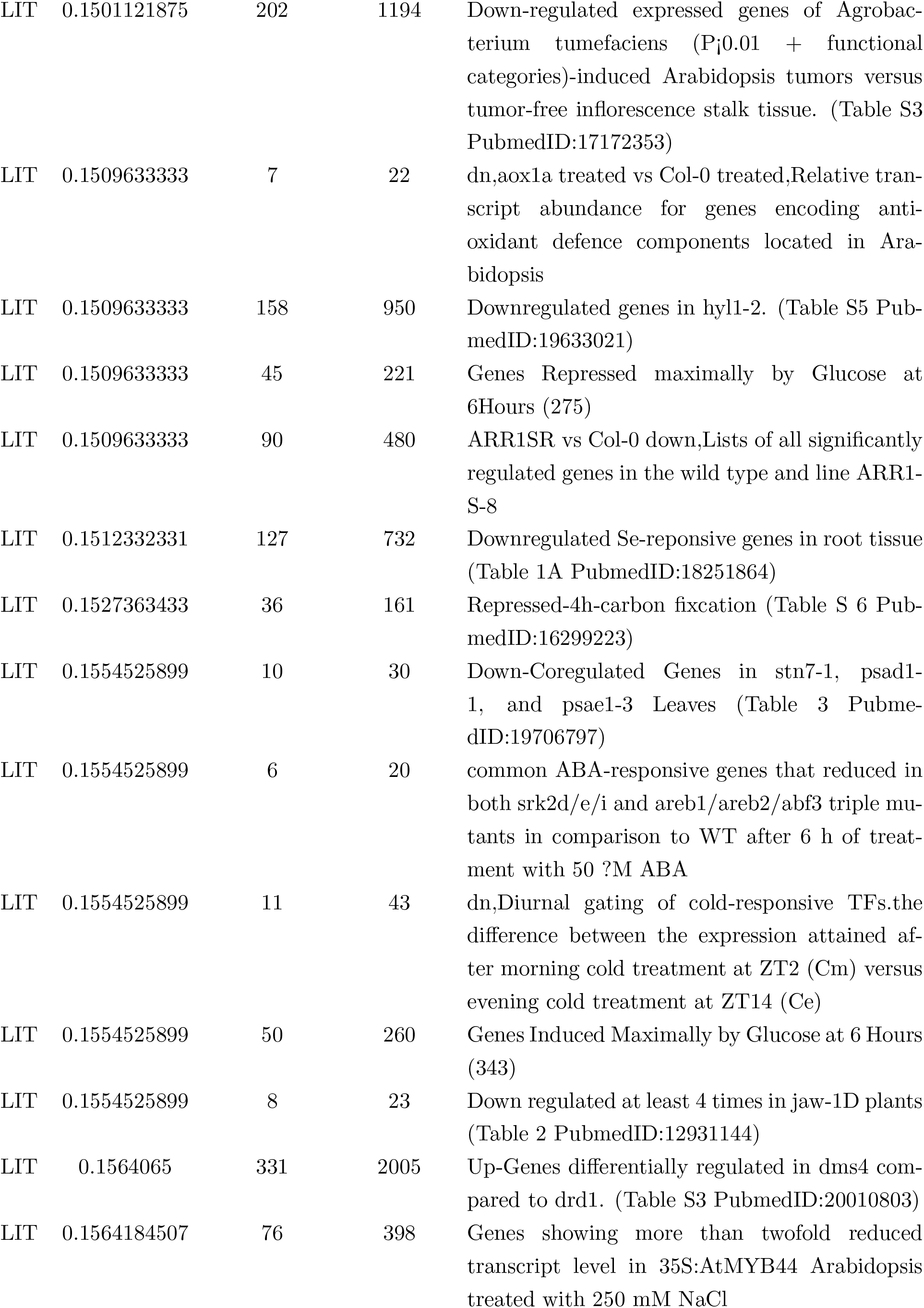
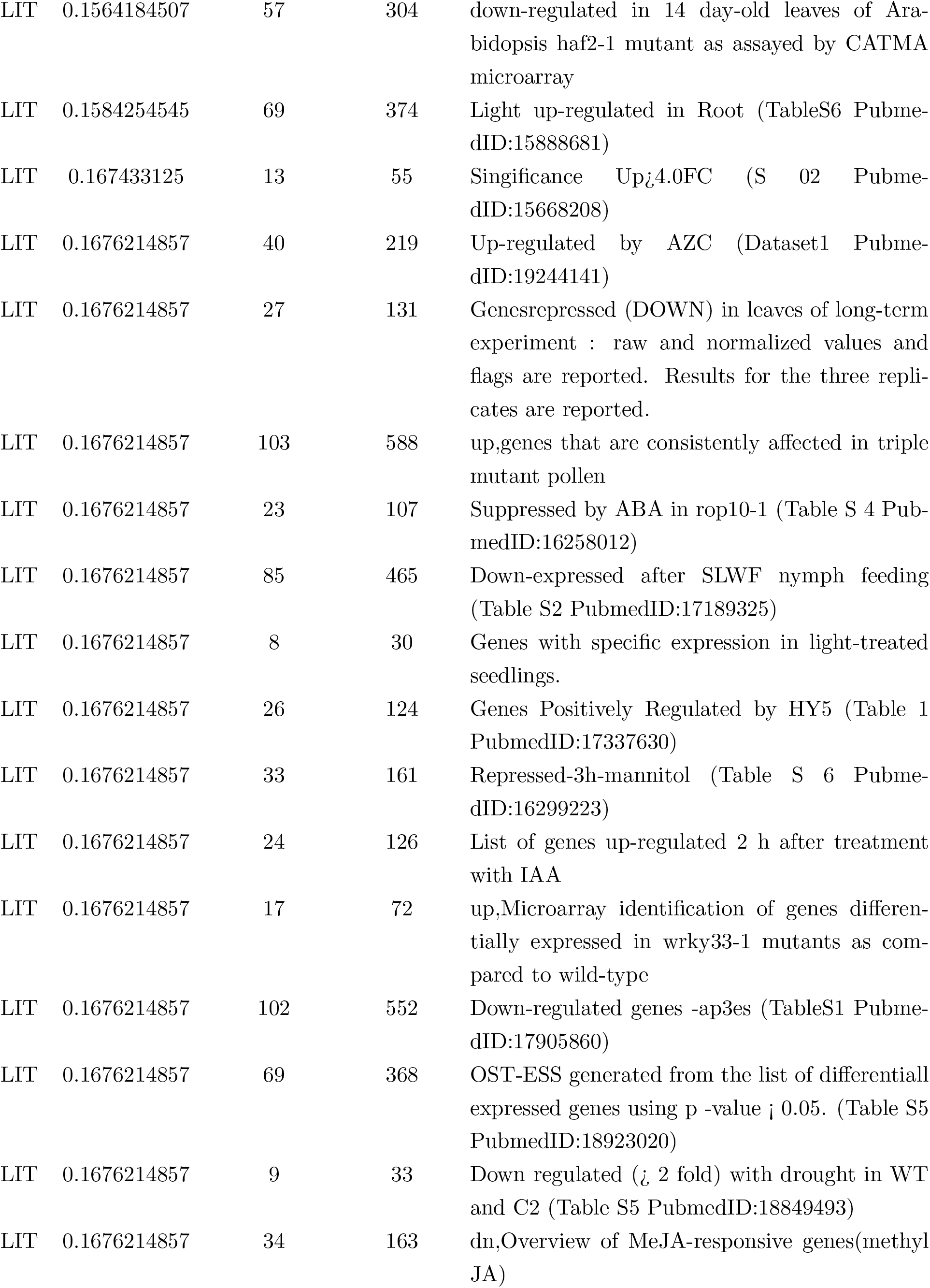
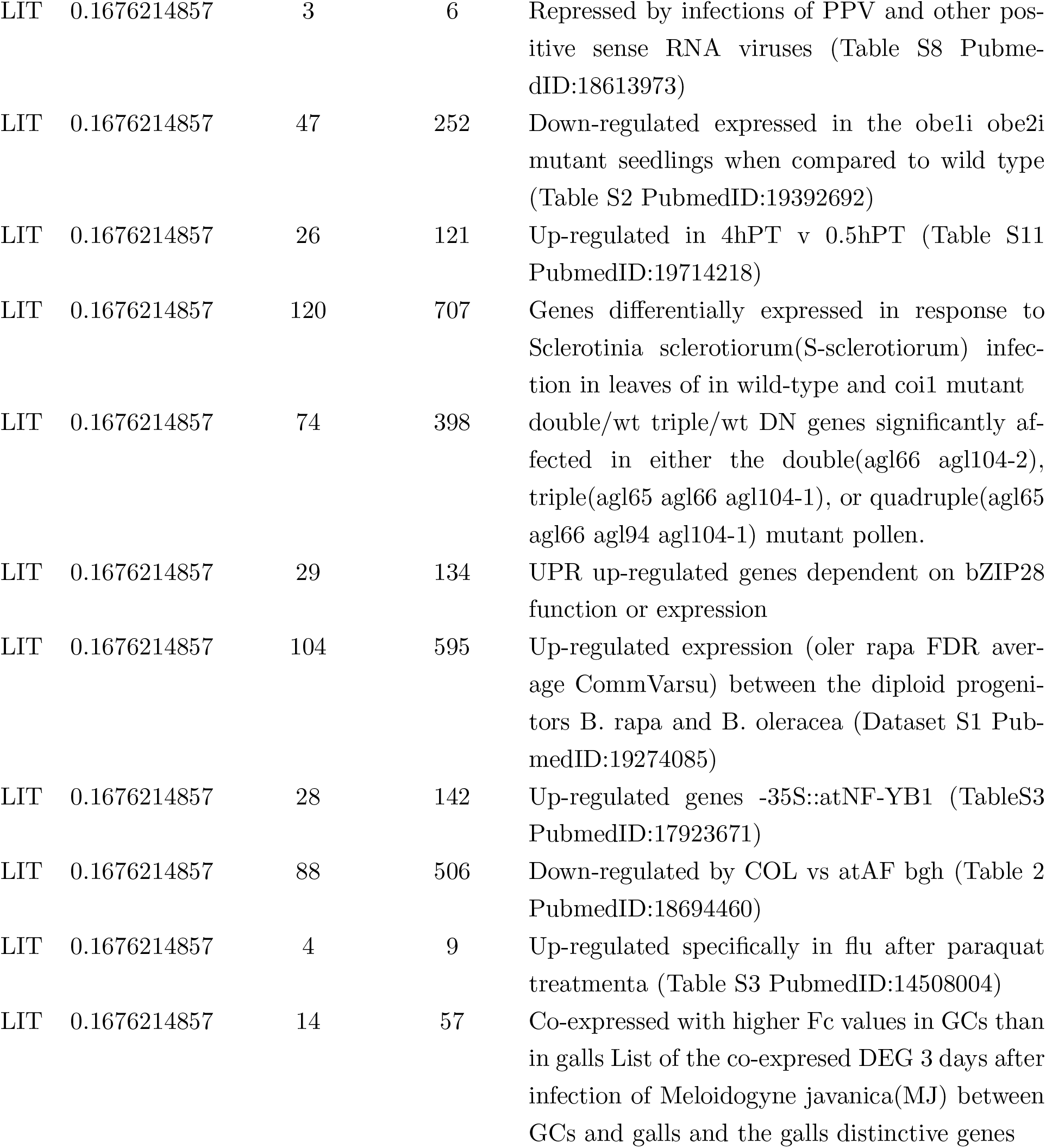
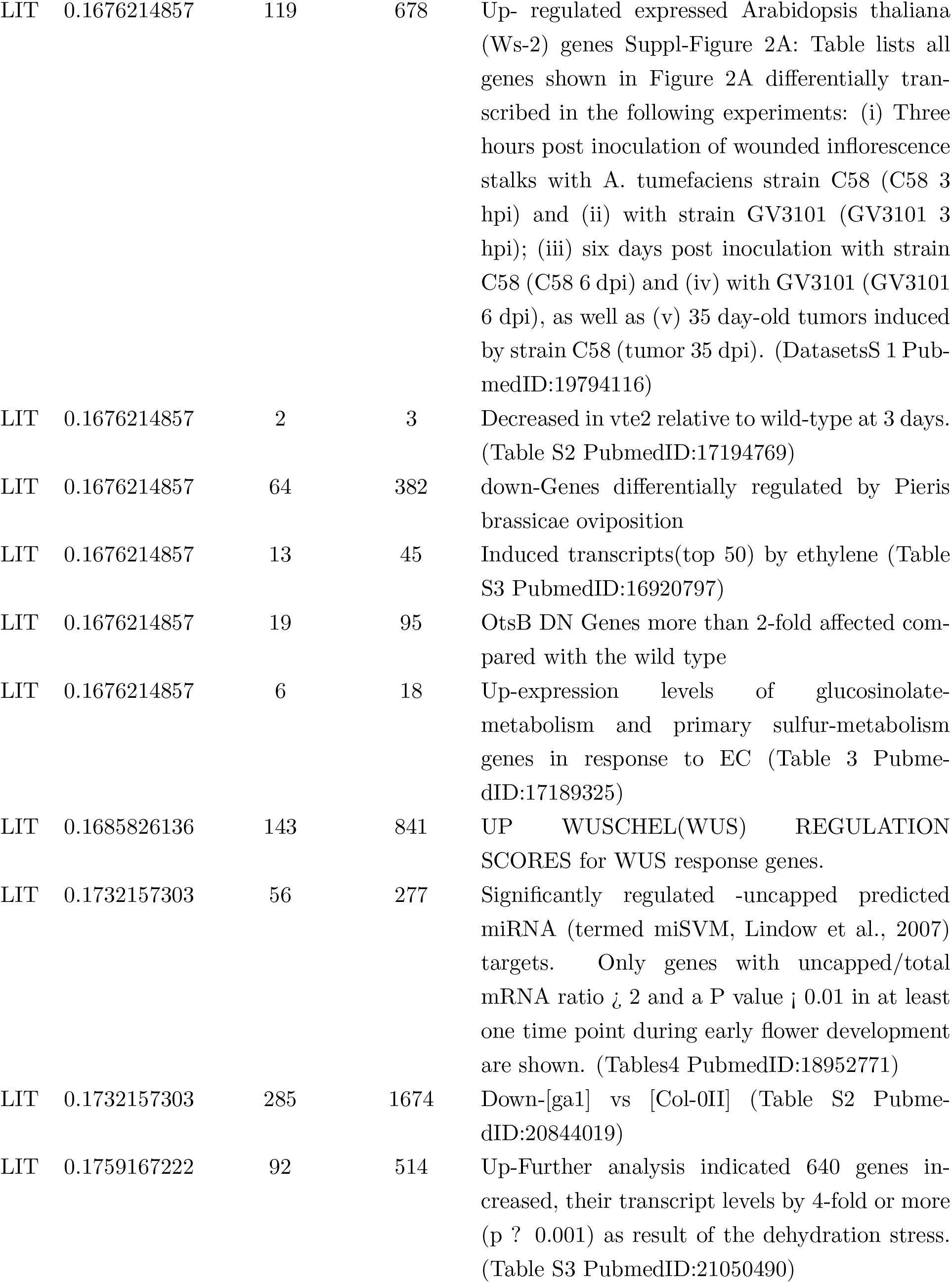
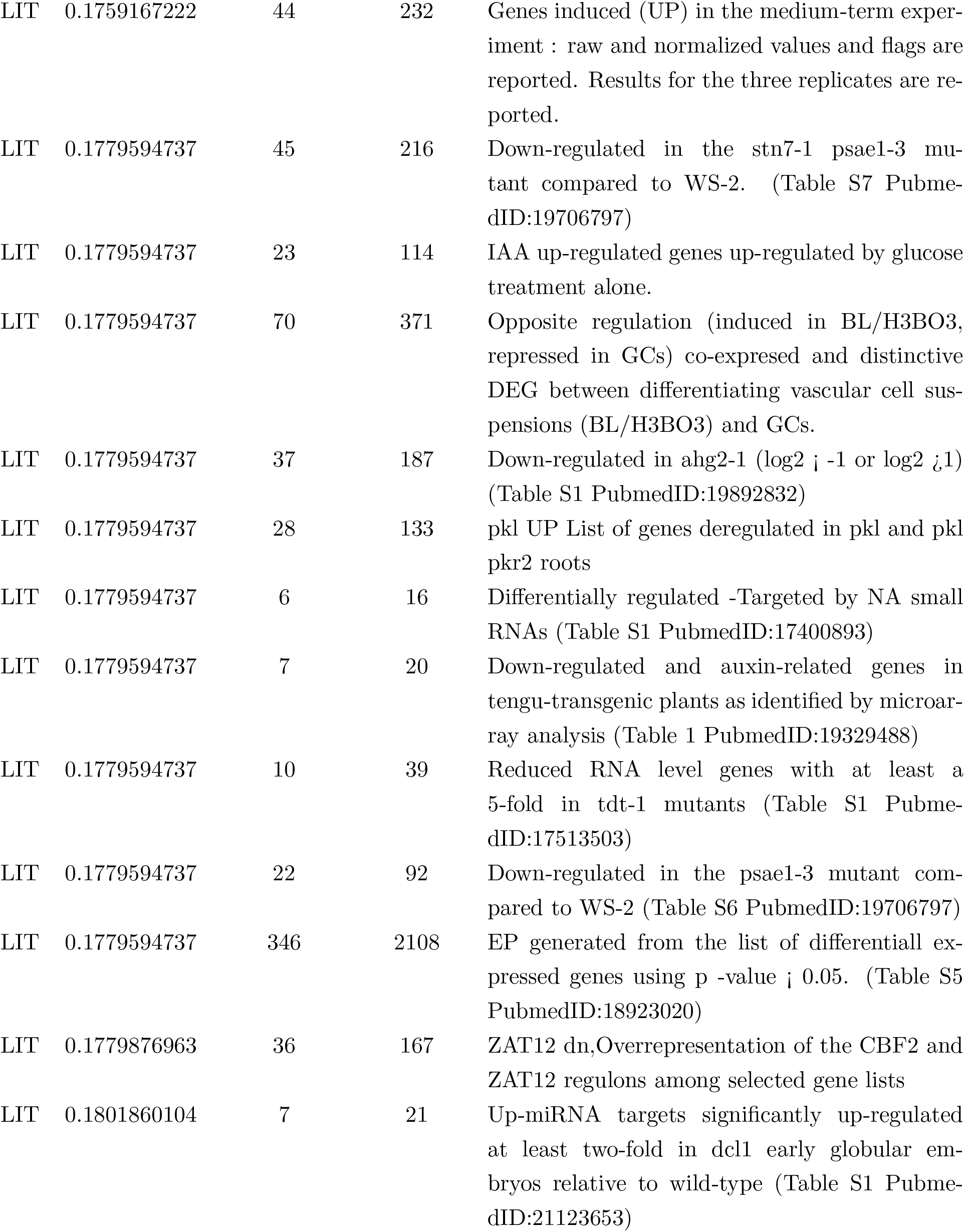
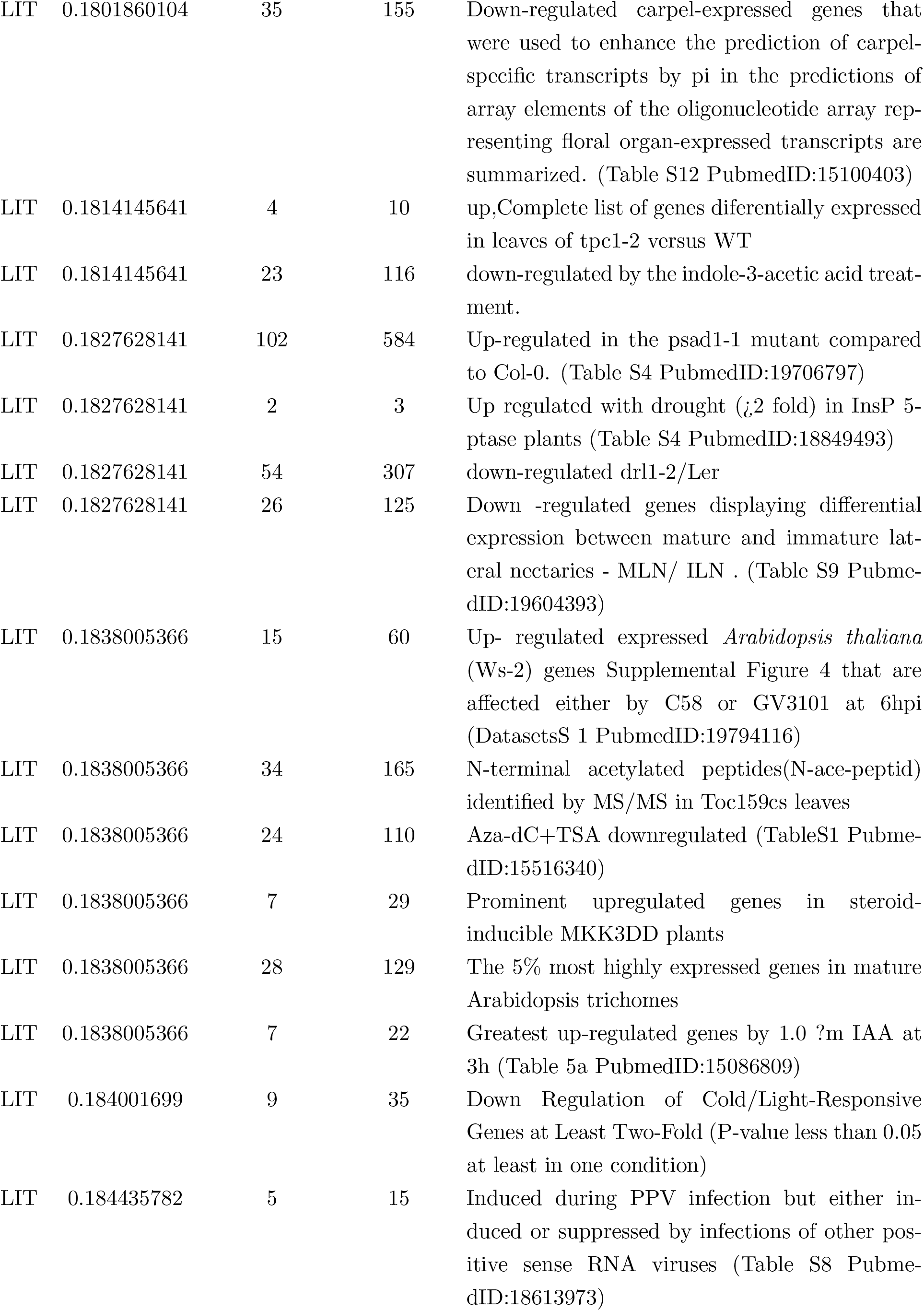
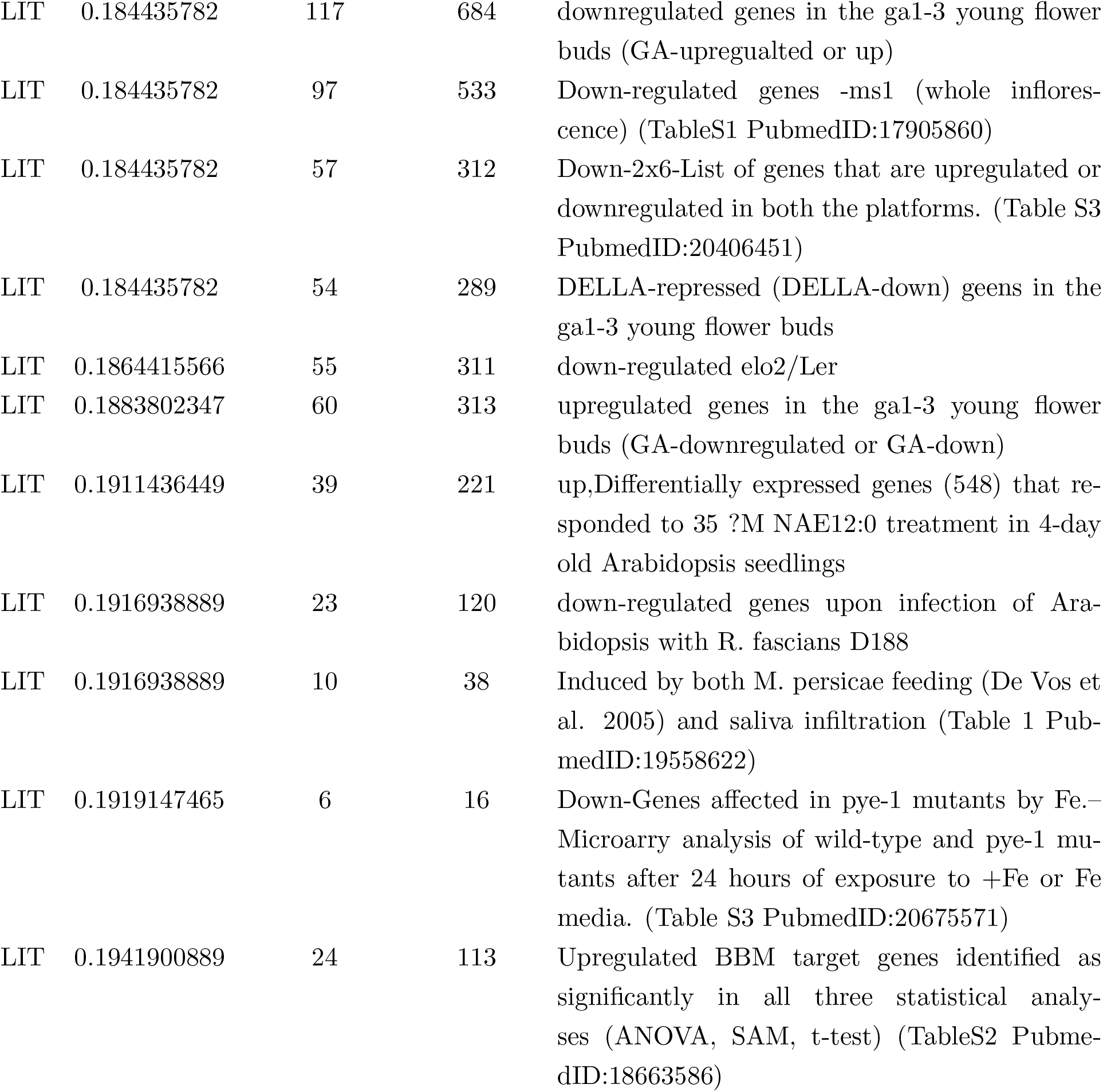
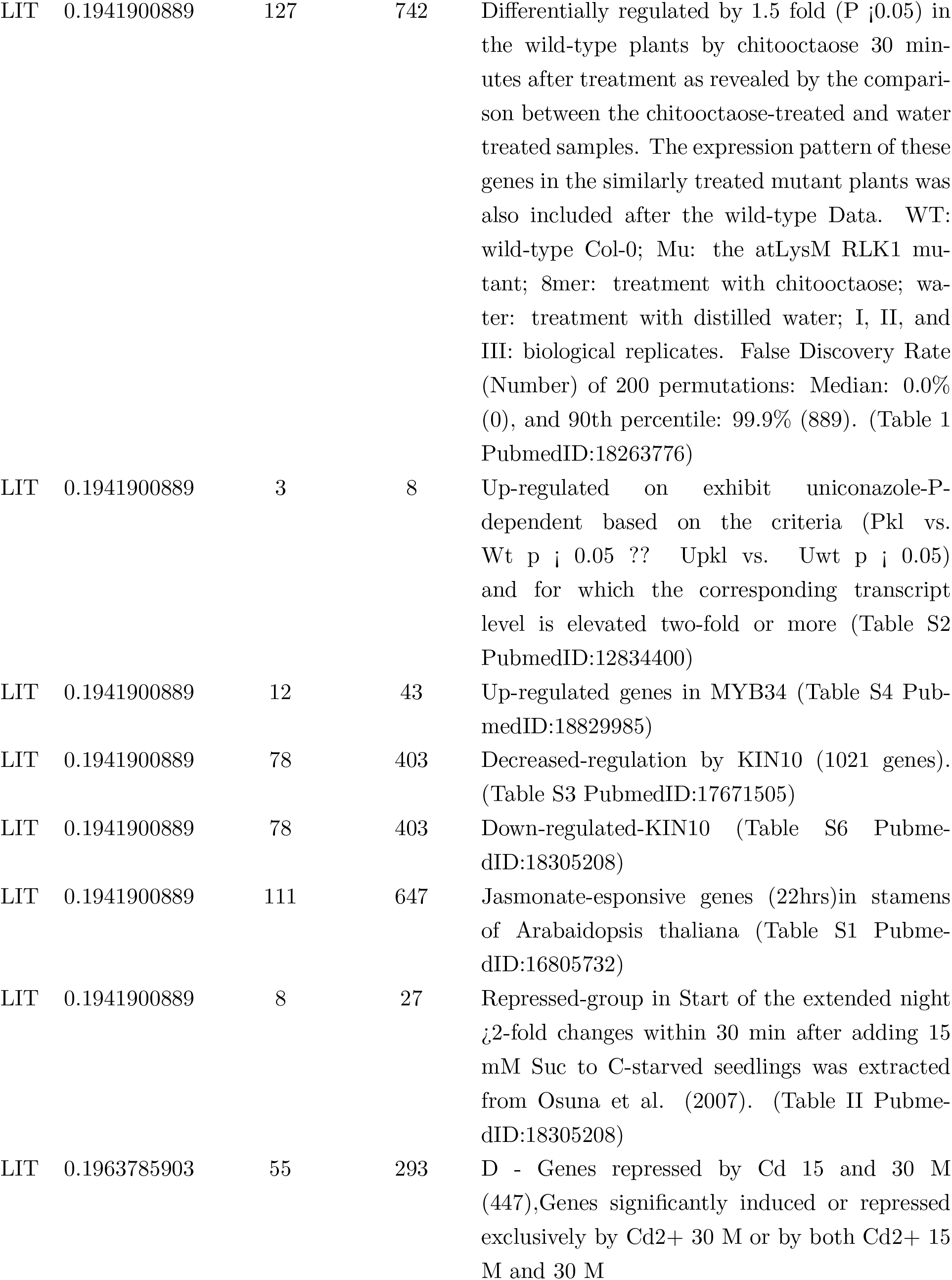
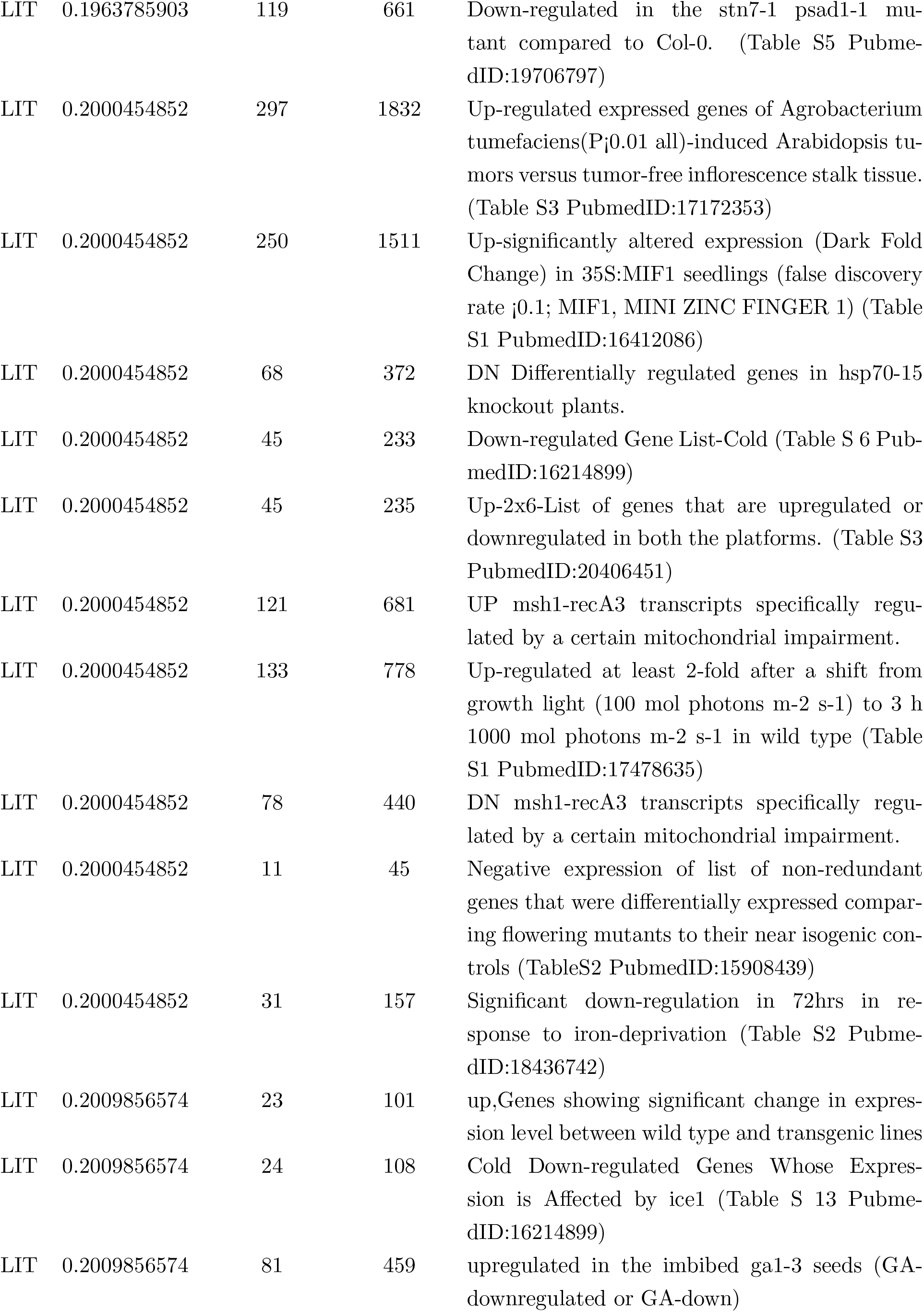
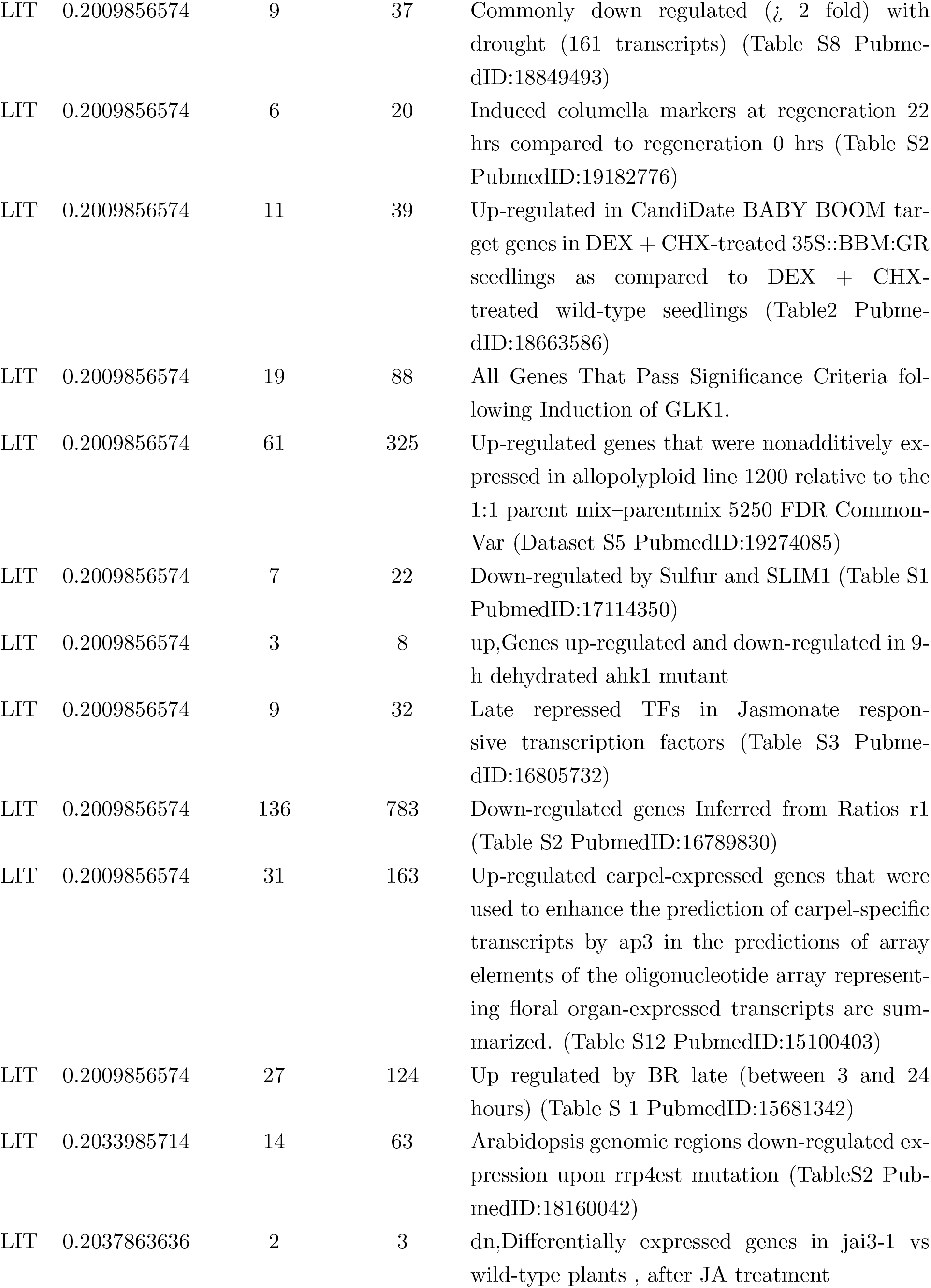
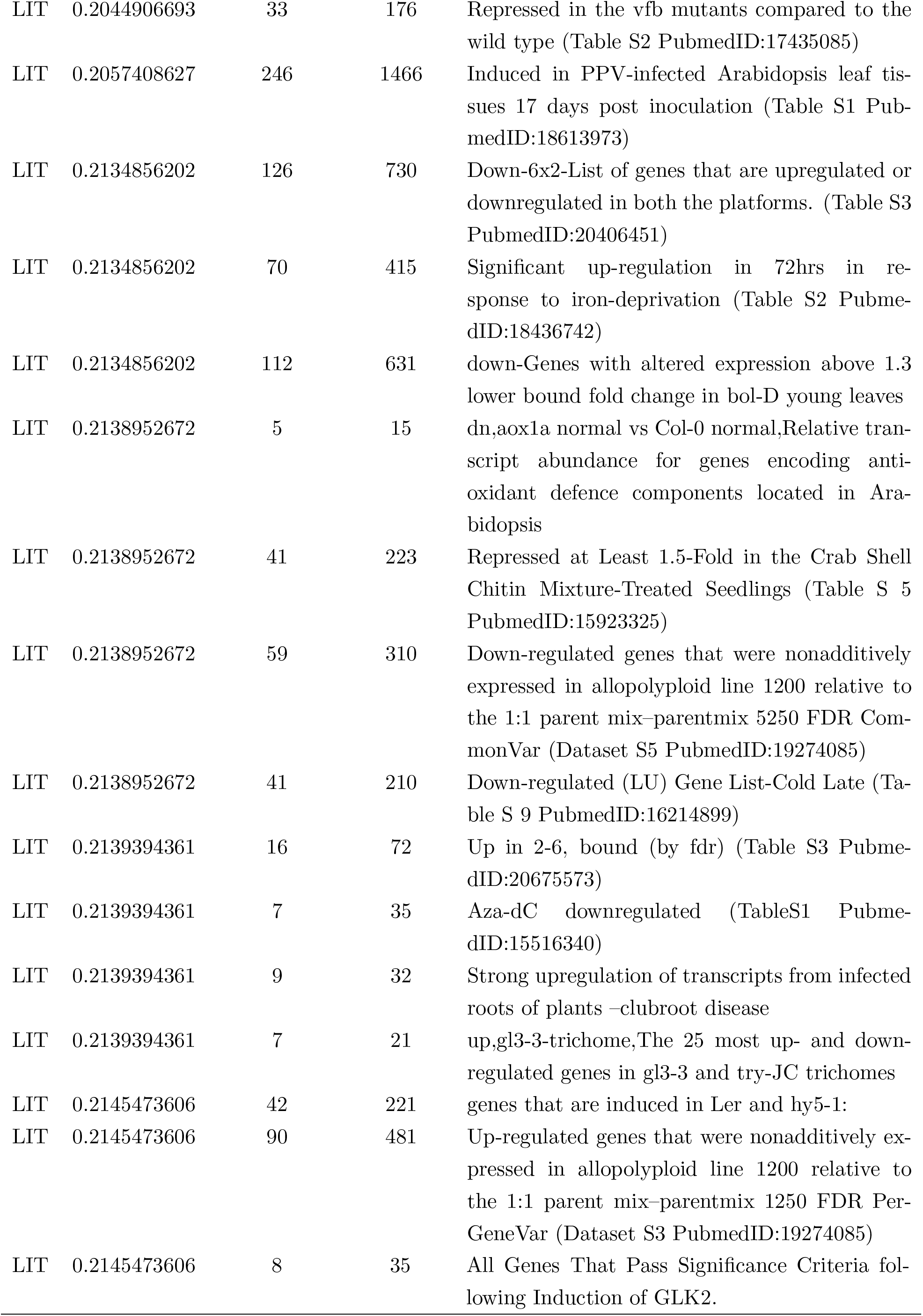
Literature collections among top *Z* signals with nominal p-value ≤ 0:05

